# Frontal Cortex Differentiates Between Free and Imposed Target Selection in Multiple-Target Search

**DOI:** 10.1101/559500

**Authors:** Eduard Ort, Johannes J. Fahrenfort, Reshanne Reeder, Stefan Pollmann, Christian N. L. Olivers

**Author notes:** Contact Information Eduard Ort, Vrije Universiteit Amsterdam, van der Boechorststraat 7, 1081 BT Amsterdam, The Netherlands. Johannes J. Fahrenfort, Vrije Universiteit Amsterdam, van der Boechorststraat 7, 1081 BT Amsterdam, The Netherlands Reshanne Reeder, Universitätsplatz 2, 39106, Magdeburg, Germany Stefan Pollmann, Universitätsplatz 2, 39106, Magdeburg, Germany Christian N. L. Olivers, Vrije Universiteit Amsterdam, van der Boechorststraat 7, 1081 BT Amsterdam, The Netherlands.

## Abstract

Cognitive control can involve proactive (preparatory) and reactive (corrective) mechanisms. Using a gaze-contingent eye tracking paradigm combined with fMRI, we investigated the involvement of these different modes of control and their underlying neural networks, when switching between different targets in multiple-target search. Participants simultaneously searched for two possible targets presented among distractors, and selected one of them. In one condition, only one of the targets was available in each display, so that the choice was imposed, and reactive control would be required. In the other condition, both targets were present, giving observers free choice over target selection, and allowing for proactive control. Switch costs emerged only when targets were imposed and not when target selection was free. We found differential levels of activity in the frontoparietal control network depending on whether target switches were free or imposed. Furthermore, we observed core regions of the default mode network to be active during target repetitions, indicating reduced control on these trials. Free and imposed switches jointly activated parietal and posterior frontal cortices, while free switches additionally activated anterior frontal cortices. These findings highlight unique contributions of proactive and reactive control during visual search.

## 1. Introduction

During search for a visual object, a mental representation of the target object is maintained in visual working memory to guide attention toward potentially task-relevant regions (Desimone & Duncan, 1995; Olivers & Eimer, 2011). In everyday situations, individuals may oftentimes try to find multiple objects at the same time, which would require the maintenance of more than one target representation. It has been shown that such multiple-target search can be challenging, often resulting in reduced search performance (Barrett & Zobay, 2014; Dombrowe, Donk, & Olivers, 2011; Found & Müller, 1996; Juola, Botella, & Palacios, 2004; Liu & Jigo, 2017; Maljkovic & Nakayama, 1994; Menneer, Barrett, Phillips, Donnelly, & Cave, 2007), raising the question as to how these multiple target representations are established for, and updated during, search – in other words, how these representations are controlled.

Recent work from one of our labs suggests that when observers look for more than one target, they may dynamically prioritize one of multiple potential target representations to guide search at any given moment (Ort, Fahrenfort, & Olivers, 2017, 2018). Specifically, we found that performance in multiple-target search depends on whether or not observers are given the opportunity to freely choose the target to select. In a gaze-contingent search paradigm, observers were instructed to always find one of two potential target colors. Importantly, they could either freely select the target to look for on a particular trial, as both targets would always be available in each search display, or the choice was imposed upon them, as only one of the two targets would be present on each trial. Eye movement latencies showed that, relative to target repeats, target switches were more costly when imposed than when made under free choice conditions. In further support of this, Van Driel, Ort, Fahrenfort, & Olivers (2019) recently conducted electroencephalography (EEG) measurements during free and imposed choice, and found that free switching between targets is associated with *pre-trial* power suppression in the beta band over midfrontal electrodes – a signal that has been linked to choice behavior (Donner, Siegel, Fries, & Engel, 2009; Spitzer & Haegens, 2017). In contrast, forced target switches elicited *post-trial* power enhancement in the delta/theta band – a signal that has been associated with conflict detection (Cavanagh & Frank, 2014; Cohen, 2014; Duprez, Gulbinaite, & Cohen, 2018). We interpret these eye movement and EEG findings within an influential framework proposed by Braver (2012), which assumes a division of cognitive control into two modes: *proactive* and *reactive* control. Proactive control is invoked and maintained in anticipation of a task, whereas reactive control is triggered whenever a conflict or unexpected event occurs. In multiple-target search, the availability of all targets in a display allows for proactive control, as observers can freely prepare for either target (cf. Arrington & Logan, 2004, 2005; Meiran, 1996). In contrast, imposing a target (i.e. by only presenting only one of the two targets in the search display) would invoke reactive control, which is reflected in a costly switching of attentional priority to the only available target. In the present study we sought to uncover the brain areas underlying free and imposed multiple-target search.

### 1.1 Brain areas involved in different modes of control

Cognitive control has been extensively investigated in the context of the implementation of, and switching between, different task sets (Meiran, 2010; E. K. Miller & Cohen, 2001). Task switches have been associated with brain regions that are considered part of a general cognitive control network that is distributed mainly over frontoparietal regions of the brain (Cole & Schneider, 2007; Dosenbach et al., 2006; Dosenbach, Fair, Cohen, Schlaggar, & Petersen, 2008; Duncan, 2010; Dove, Pollmann, Schubert, Wiggins, & Yves Von Cramon, 2000; Fedorenko, Duncan, & Kanwisher, 2013; Kim, Cilles, Johnson, & Gold, 2012; Liston, Matalon, Hare, Davidson, & Casey, 2006; Power & Petersen, 2013, A. B. Smith, Taylor, Brammer, & Rubia, 2004). However, it is unknown whether similar brain areas are also involved in switching representations within one and the same task, which is the case when observers hold multiple target representations for the same visual search task, and how this would differ for circumstances that enable different modes of control.

The distinction between proactive and reactive control has mostly been studied in the context of interference control across various domains, such as interference between competing working memory representations (Burgess & Braver, 2010; Marklund & Persson, 2012), between competing visual stimuli (De Pisapia & Braver, 2006; Jiang, Beck, Heller, & Egner, 2015), or between competing stimulus-response mappings (Braver, Reynolds, & Donaldson, 2003; Jiang, Wagner, & Egner, 2018; Ryman et al., 2018; Sohn, Ursu, Anderson, Stenger, & Carter, 2000). The mode of control is commonly induced by manipulating the likelihood (or predictability) of upcoming interference, following the assumption that whenever individuals anticipate interference, they will strengthen proactive control. Some studies suggest that proactive and reactive control are governed by different brain areas, but the findings are somewhat inconsistent, which may be related to the different paradigms used, and to an emphasis on differences in the temporal dynamics (with proactive assumed to occur prior to task onset, while reactive follows conflicting events). Based on their reviews of the literature, Braver (2012) as well as Irlbacher, Kraft, Kehrer, & Brandt (2014) have suggested that both modes of control are governed by a similar set of brain areas, but might be activated with different dynamics, as proactive control can be instantiated in advance. These areas include the lateral prefrontal cortex and posterior parietal cortex, with relatively minor differences between them, whereas reactive control may additionally recruit midfrontal regions when there is conflict detection involved. Similar brain areas may therefore be involved during control over target selection in multiple-target search.

Target selection in multiple-target search is associated with shifts in feature-based attention between target-defining features. Such shifts of feature-based attention have previously been linked to activity primarily in posterior parietal (i.e. superior parietal lobule) and posterior, lateral frontal regions (i.e. inferior frontal junction and dorsal premotor cortex; Greenberg, Esterman, Wilson, Serences, & Yantis, 2010; Liu, Slotnick, Serences, & Yantis, 2003; Pollmann, Weidner, Müller, Maertens, & Von Cramon, 2006; Pollmann, Weidner, Müller, & Von Cramon, 2000; Slagter et al., 2006, 2007; Wager, Jonides, & Reading, 2004). However, in most of these studies, a feature shift was also associated with a change in response (Greenberg et al., 2010; Liu et al., 2003; Pollmann et al., 2006; Slagter et al., 2006, 2007). Moreover, these studies did not directly compare different modes of control over such shifts. They measured activity in response to either cue- or task-induced changes of the task-relevant feature, but did not juxtapose self-generated (free) to stimulus-induced (imposed) changes.

In one recent study, Gmeindl et al. (2016) did compare cue-induced to self-generated (i.e. freely chosen) shifts of spatial attention. They found similar posterior parietal activity during both types of shifts, while self-generated shifts additionally activated the medial frontal cortex and lateral frontopolar cortex. These medial frontal and frontopolar regions have also previously been shown to be related to voluntary versus imposed action selection (Demanet, De Baene, Arrington, & Brass, 2013; Forstmann, Brass, Koch, & von Cramon, 2006; Orr & Banich, 2013; Passingham, Bengtsson, & Lau, 2010; Soon, Brass, Heinze, & Haynes, 2008; Taylor, Rushworth, & Nobre, 2008; Wisniewski, Goschke, & Haynes, 2016; Wisniewski, Reverberi, Tusche, & Haynes, 2015; J. Zhang, Kriegeskorte, Carlin, & Rowe, 2013), and have been argued to be involved in the evaluation of alternative goals in the context of exploratory behavior (Mansouri, Koechlin, Rosa, & Buckley, 2017; Pollmann, 2016). The same areas may therefore be involved when observers choose to change target in visual search, but this is currently unknown.

### 1.2 The present study

We sought to investigate differences in the locus or level of activated brain regions when proactive and reactive control mechanisms operate in a context of multiple-target search. Specifically, we set out to test whether differences between free and imposed switches between targets during visual search for multiple objects are accompanied by differences in brain activity that might be linked to proactive and reactive control processes. To that end, we adopted a fast-paced, gaze-contingent eye tracking paradigm (Ort et al., 2017; illustrated in Figure 1A) in combination with event-related fMRI. Participants were always instructed to look for two color-defined targets and to make an eye movement towards one of them on every trial. In one block type, both potential targets were present in a search display and participants were free to select either of them. In the other type of block, only one target was present and the choice was imposed. We instructed participants to either make (when choice was free) or expect (when choice was imposed) target switches. We reasoned that free target switches would be associated with proactive, preparatory control mechanisms, while imposed switches would result in reactive, compensatory control mechanisms. Differential neural activity for each switch type would constitute evidence for proactive and reactive control having unique contributions to target selection during multiple-target search. Based on the literature on both task and attention shifts, we expected to find switch-related activity in the posterior parietal and posterior frontal cortex for both switch types. In addition, we were interested to explore potential differences in the control network for free and imposed target switches. In line with the literature on self-generated versus externally-cued choice, we expected activity in the lateral frontopolar cortex as well as the medial frontal cortex to be selectively active during free switches.

**Figure 1.**
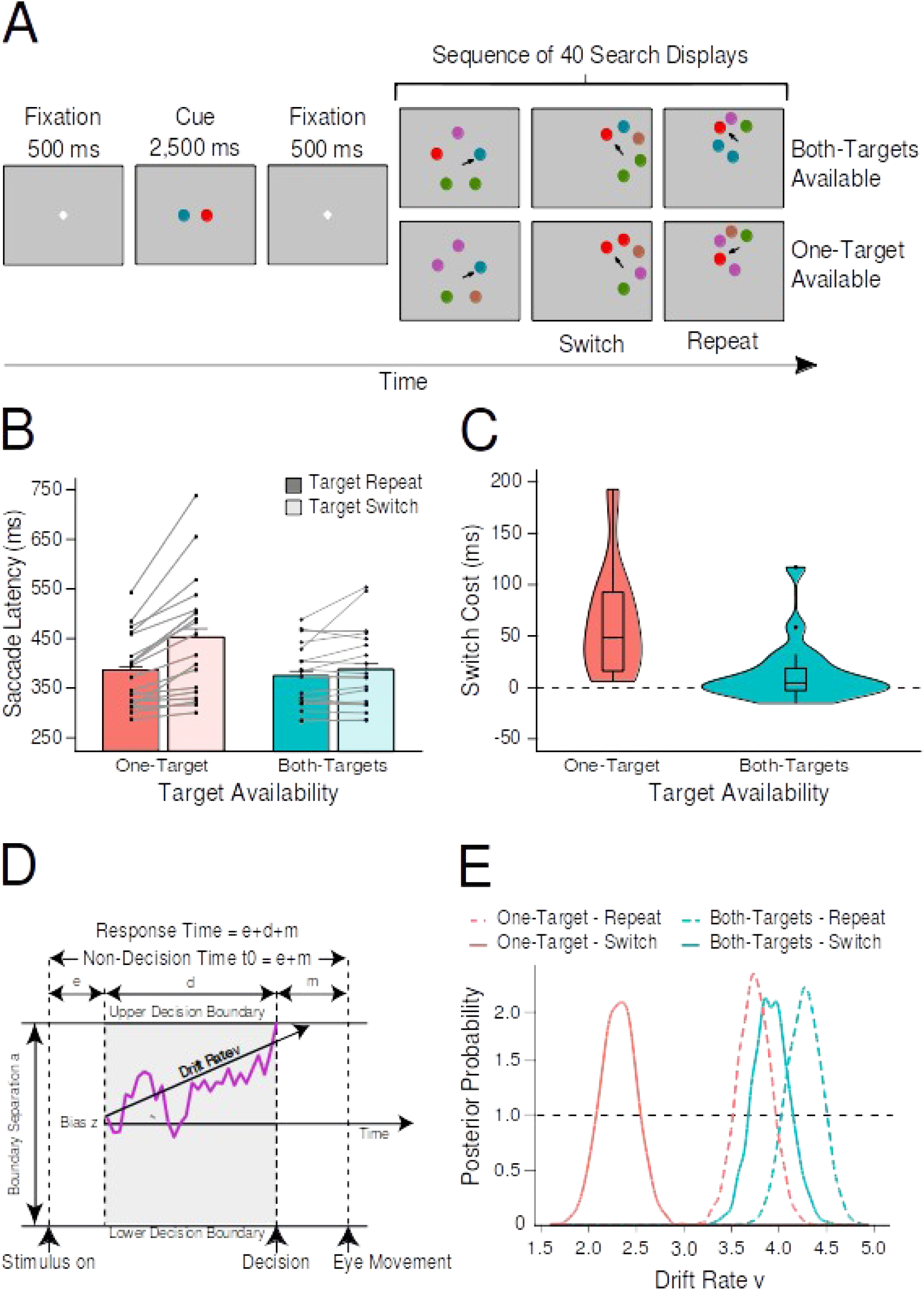
Illustration of the study design and behavioral results. A) Each block began with a cue indicating the two target colors for the subsequent sequence of 40 search displays. Depending on the target availability condition, each search display contained either one of the two target colors or both of them. In case of only one target color being available, there could still be two targets carrying that color, to equate for the mere number of targets present (see section 2.3). Participants were required to fixate one of the targets (indicated here by an arrow, which was not present in the display); this triggered the next display, which appeared on an imaginary annulus surrounding the location of the previously fixated target. B) The bar plots represent the mean saccade latencies on switch trials and repeat trials for each level of target availability (one target vs. both targets). The gray lines represent the mean saccade latencies for each observer individually. Error bars represent the upper limit of the within-subjects 95% confidence intervals (Morey, 2008). C) The violin plots depict the distribution of switch costs, which were computed by subtracting repeat saccade latencies from switch saccade latencies, separately for the target availability conditions. The horizontal lines in the box plots represent the first, second (median), and third quartiles. The vertical lines represent the distance between minimum (lower quartile − 1.5 * interquartile range) and maximum (upper quartile + 1.5 * interquartile range). Single dots indicate individual outliers. D) Schematic and simplified equation of a drift diffusion model (adapted from Kloosterman, et al., 2019). *e* denotes encoding time, *d* denotes decision time, and *m* denotes motor execution. E) HDDM results indicating the posterior probability distributions for drift rates, separately for all experimental conditions.

## 2. Methods

### 2.1. Data and code availability

Data and code was made publicly available on *osf.io* (https://osf.io/a8vxn). Unthresholded statistical maps were uploaded to *neurovault.org* (https://neurovault.org/collections/5550/).

### 2.2 Participants

A sample of 22 participants (age: 21-35 years, M = 27.3; 10 females, 12 males) was recruited from the subject pool of the Leibniz Institute of Neurobiology in Magdeburg. Three individuals were excluded due to insufficient eye tracking accuracy, reducing the final sample to 19 participants. All participants gave written consent according to the Declaration of Helsinki and were reimbursed with 30 Euros. They reported normal or corrected-to-normal visual acuity and color vision and were naive to the purpose of the experiment. The study was approved by the research ethics board of Otto-von-Guericke University Magdeburg.

### 2.3 Stimuli, procedure, and design

The stimulus set consisted of five colored disks with a radius of 0.6 degrees visual angle (dva). These colors were blue (RGB-values: 0, 130, 150), red (240, 0, 0), green (70, 135, 0), brown (175, 100, 75), and purple (180, 80, 170). All stimulus colors were isoluminant (M = 21 cd/m^2^) and presented on a uniform gray background (197, 197, 197). A search display was composed of five disks placed on an imaginary annulus around fixation with a radius randomly drawn from values between 3.6 and 4.4 dva around the starting point. Any two adjacent disks had an angular distance of at least 45 degrees.

A block was initiated once participants steadily fixated a central white dot. First, a white fixation cross was presented in the center of the screen for 500 ms, followed by the cue display for 2,500 ms and another fixation cross for 500 ms (Fig. 1A). In the cue display, two colored disks were presented 1.0 dva to the left and right of fixation to mark these colors task-relevant for the upcoming sequence of 40 search displays. In each search display, participants were required to select a target-colored disk among a set of five disks by making a saccade toward it. After target fixation, the search display disappeared and the fixated target was replaced by a black ring to provide participants with a fixation point during the intertrial interval (uniformly jittered between 950 to 1050 ms). Because the coordinates of the previously fixated target served as the starting point for the next display, the search displays moved across the screen throughout a block. To make sure that search displays would fall within the margins of the screen, stimuli were moved closer to each other on that part of the annulus that was closest to the center of the screen, whenever a search display approached an edge of the screen. Importantly, the relative positions of the targets in the search display remained unpredictable. Saccades had to land within a radius of 0.9 dva to the center of a target to trigger the next search display. If participants fixated one of the distractors, they received auditory error feedback and were required to make a corrective eye movement toward a target. The search was aborted if no target was fixated within 3,000 ms, and a new search display appeared, centered at the same location.

There were two main factors in this experiment: *target availability* (whether only one or both targets were present in the search display), and *transition type* (whether target selection switched or repeated from one trial to the next). The target availability factor was controlled at the block level by presenting either only one, or both of the targets in the display. In the *both-targets condition*, both cued targets appeared in the search display. In the *one-target condition*, only one of the targets was present. The transition type factor (*target repeat* vs. *target switch*) was determined by the observer (both-targets blocks) or by a random sampling procedure (one-target blocks).

In both-targets blocks, participants were instructed that they were free to fixate either of the target colors. However, to make sure that there would be sufficient switch trials, and that the time between two consecutive switch trials would be long enough for the BOLD response triggered by each of them to not overlap, it was emphasized to participants that the total number of target switches in a block of 40 search displays should roughly be in the range of five to eight. The sampling procedure in one-target blocks then randomly selected (with replacement) a sequence of repeat and switch trials from a pool of sequences that were recorded during both-targets blocks. The motivation behind this was to match one-target blocks and both-targets blocks in terms of switch rate and number of consecutive repeat trials. Importantly, neither features nor positions of the stimuli were replayed but only the sequence of switch and repeat trials. Because we did not yet have any sequences to present at the outset of the experiment, we initialized a pool with four arbitrarily prespecified sequences of switch and repeat trials (one each for five, six, seven, and eight switches per block). To check whether switch rates indeed did not differ between target availability conditions, we ran a paired-samples *t*-test and found a slight, but non-significant difference (both-targets available: 6.4 switches, one-target available: 6.9 switches, *t*(18) = 1.9, *p* = .07, *Cohen’s d* = 0.49, *BF_10_* = 1).

Both-targets available and one-target available blocks would differ not only in terms of target availability, but also in the mere number of targets in the display, which would make the one-target available condition more difficult than the both-targets available condition. Therefore, we included trials in the one-target available condition in which there were two target objects, but both carried the same target color, so that still only one target color was present in the search display (*target duplicate*, e.g. on blocks in which red and blue were task-relevant, there could be trials with two red targets or two blue targets, but never with a red and a blue target). In addition, we included trials in which two distractors shared a color (*distractor duplicate*; e.g., on blocks in which red and blue were task-relevant, there could be trials with two green items, but only one red or one blue item), so that participants could not identify the target object simply by detecting a feature duplicate. Likewise, both-targets blocks also contained target duplicate trials (two out of three targets had the same color; e.g., on blocks in which red and blue were task-relevant, we had trials with one red and two blue target objects) as well as distractor duplicate trials (e.g., on blocks in which red and blue were task-relevant, there were trials with one red target, one blue target, and two green distractors). As a result, in each target availability condition, one half of trials contained a target duplicate and the other half contained a distractor duplicate. This way, neither the number of targets nor the number of unique colors in the display was predictive of target availability. Supplementary Table S1 provides schematic representations of all types of search displays. Past experiments using a similar paradigm have confirmed that behavior and ensuing switch costs are consistently unaffected by this manipulation (Ort et al., 2017, 2018), as we also confirm here (see section 3.1). Furthermore, to investigate whether there would be other experimental variables that influenced the choice behavior of the participants we ran a series of control analyses. These analyses are summarized in the Supplementary Material online and Figure S1.

Because we did not have an eye-tracker available outside of the scanner, participants practiced a version of the task in which target selection responses were made using mouse tracking instead of eye movements (although this naturally involves making an eye movement to the target too). They performed this task before the fMRI session started until they felt confident they understood the task structure. A scanning session consisted of nine functional runs, each seven minutes long. One participant requested to terminate the last run early (leaving eight runs of data), while another participant completed ten runs because he expressed the wish to do another run as he liked doing the experiment (data of this run were included). In a single run, both-targets and one-target blocks alternated repeatedly until the end. For the first block in a run, the target availability condition switched relative to the last complete block of the previous run. To make sure that both block types would occur each at least twice per run and that a block would not exceed the run duration, a block was interrupted after 88 seconds (mean complete block duration = 73 s), or five seconds before the run finished. This resulted in up to five complete blocks per run and, on average, 32 complete blocks per session. Nevertheless, incomplete blocks were still analyzed up to the point of termination.

### 2.4 Apparatus and functional MRI acquisition

The experiment was designed and presented using the OpenSesame software package (version 3.1.9; Mathôt, Schreij, & Theeuwes, 2012) in combination with PyGaze (version 0.6), an eye-tracking toolbox (Dalmaijer, Mathôt, & Van der Stigchel, 2013). Stimuli were back-projected on a screen with a resolution of 1,280 x 1,024 pixels, at a viewing distance of 60 cm. Participants viewed the screen via an IR-reflecting first surface mirror attached to the head coil. Eye movements were recorded with the EyeLink 1000 remote eye-tracking system, (SR Research, Mississauga, Ontario, Canada) at a sampling rate of 1000 Hz. The experimenter received real-time feedback on system accuracy on a second monitor located in an adjacent room. After every run, eye-tracker accuracy was assessed and improved as needed by applying a 9-point calibration and validation procedure (mean calibration error was 0.48 dva).

Images were acquired using a 3 Tesla Philips Achieva dStream MRI scanner with a 32 channel head coil. Functional images were recorded using a T2*-weighted single-shot gradient echo-planar images sequence with following parameters: 35 axial slices parallel to the AC-PC axis (ascending order), slice thickness = 3 mm, in-plane resolution = 80 x 78 voxels (3 mm x 3 mm), FOV = 240 mm x 240 mm, inter-slice gap of 10% (0.3 mm), whole-brain coverage, TR = 2 s, TE = 30 ms, flip angle = 90°, parallel acquisition with sensitivity encoding (SENSE) with reduction factor 2. After the first five scans were discarded, 210 scans were acquired per functional run. Structural images were recorded using a T1-weighted (T1w) MPRAGE sequence with following parameters: 192 slices, slice thickness = 1 mm, in-plane resolution = 256 x 256 voxels (1 mm x 1 mm), FOV = 256 mm x 256 mm, TR = 9.7 ms, TE = 4.7 ms, inversion time = 900 ms, flip angle = 8°. Distortions of the B0 magnetic field, as well as pulse oximetry and respiratory trace were recorded, but these data were not further processed.

### 2.5 Eye-tracking data preprocessing

We compared the saccade latencies of eye movements (dwell time before a saccade was executed) for repeat trials (current target category the same as the previous one) with those for switch trials (current target category different from the previous one) for both target availability conditions separately. We took the first saccade after search display onset with an amplitude threshold of 1 dva around initial fixation, provided that a saccade was directed toward the selected target (i.e. its direction deviated less than 30 angular degrees from the vector from fixation to the target). This resulted in an average of 25.5% of all trials being removed. Note, as we used only the first saccade of a trial and participants needed to select a target before actually fixating it, our paradigm measures covert selection process. Next, a saccade latency filter was applied, in which saccades quicker than 100 ms and slower than 3 standard deviations above the block mean for that participant were excluded (2.2% of all search displays). If no target was being fixated, as could have happened when the eye tracker calibration had deteriorated too much for that trial, both the current as well as the next search display were excluded because neither could be labeled as a switch or repeat (10.6% of all search displays). For the same reason, we excluded the first search display of each block (2.7% of all search displays). If the distance between the stimuli was lowered to prevent the search displays to fall outside the screen, two stimuli could be too close to each other to unambiguously decide which of the two was fixated. Trials on which this happened were also excluded (7.4% of all search displays). In total, 34.4% of all trials were thus removed during preprocessing (note that a single trial could meet multiple exclusion criteria). This is a typical rejection rate for this paradigm (Ort et al., 2017, 2018, van Driel et al., 2019). Inferential statistics were carried out with the *afex* R-package (Singmann et al., 2016).

### 2.6 Hierarchical drift diffusion modeling

To gain more insight into target selection beyond simple comparisons of mean saccade latencies across conditions, we also performed drift diffusion modeling (DDM) on our data. DDMs can estimate latent decision-related parameters in two-alternative choice experiments based on response time distributions and choice probabilities (Wiecki, Sofer, & Frank, 2013). For example, participants might be more cautious to respond when only one target is available than when both target colors are present. Similarly, participants might need longer to select a stimulus to fixate on switch trials when only one target is present. In these models, decision-making is assumed to be a noisy information accumulation process in favor of one or the other alternative that continues until a threshold for one option is exceeded and the corresponding response is executed (e.g. Ratcliff & Rouder, 1998). We used a hierarchical Bayesian DDM, as implemented in the python-library *HDDM* (version 0.6; Wiecki et al., 2013), which has the advantage of simultaneously estimating group and individual-subject parameters as well as obtaining a measure for the estimates’ uncertainty.

The standard DDM framework provides estimates for four parameters: drift rate *v*, non-decision time *t*, boundary separation *a,* and starting point *z* (see Figure 1D). The drift rate represents the speed with which evidence is accumulated during a decision process. It is related to the difficulty of a decision, with hard decisions corresponding to low drift rates and easy decisions to high drift rates. The non-decision time signifies the time needed to encode the stimulus and execute the motor response and is therefore not related to the decision process itself. The boundary separation parameter reflects how much evidence needs to be accumulated before a decision is made, therefore representing response caution (speed-accuracy trade-off). Close boundaries lead to quick and more inaccurate decisions, whereas wide boundaries lead to slower, but more accurate decisions. The starting point denotes whether there is an a priori bias for one of the options.

It has been shown that task repeats are associated with higher drift rates than target switches, plausibly reflecting faster evidence accumulation when a target is repeated (e.g. Karayanidis et al., 2009; Schmitz & Voss, 2012). Based on this finding, we also expected to find higher drift rates for target repeats than for target switches. However, if individuals prepare a target prior to search display onset in both-targets blocks, drift rates for free target switches should likely be higher than for imposed switches, potentially even be as high as the drift rate for target repeats. Furthermore, in Ort and colleagues (2017), we found base rate differences between both-targets and one-target blocks in terms of saccade latencies: Saccades were generally faster in both-targets blocks than in one-target blocks, irrespective of transition type. This effect might imply strategic differences, such as increased response caution when only one target was available relative to when both were present. To investigate this hypothesis, we estimated a separate boundary separation for both-targets and one-target blocks. As the boundary separation is usually assumed to be under control of individuals and switch and repeat trials were unpredictable in one-target blocks, we did not separately estimate this parameter for repeat and switch trials.

Unlike in standard two-forced choice tasks, there was no single correct or incorrect response in our task. In fact, on every trial, participants could make five possible responses, corresponding to each stimulus (targets and distractors). Therefore, to make our paradigm compatible with the DDM framework, we did not consider individual stimuli as response options, but only distinguished correct (saccades to targets) from incorrect (saccades to distractors) responses. To test our hypotheses about the influence of the experimental conditions on drift rate and boundary separation, we ran four models, in which we manipulated which parameters were free to vary across experimental conditions. These models were: (1) basic model, in which both boundary separation (*a*) and drift rate (*v*) were fixed across conditions; (2) decision boundary model, in which *a* could vary between target availability conditions and *v* was fixed; (3) drift-rate model, in which *v* could vary between target availability and transition type conditions, and *a* was fixed; (4) full model, in which *a and v* could vary between target availability conditions, and *v* could also vary between transition type condition. We did not estimate intertrial variability of starting point and drift rate in any of the models and fixed the estimate for starting point and non-decision time across conditions, as condition-specific differences in those parameters were implausible. Furthermore, we chose to use informative priors (see Wiecki et al., 2013). Nevertheless, once we identified the best model, we also ran it with non-informative priors as control analysis; the results were virtually identical. Supplementary Table S2 includes the full specifications of all models that were tested.

For every model, 50,000 steps were sampled with Markov Chain Monte Carlo (MCMC). The first 20,000 samples were discarded (“burn in”) and only every fifth sample was kept (“thinning”) to facilitate convergence. Convergence was tested by visually inspecting all posterior distributions (mc-trace, auto-correlation and marginal posterior histogram) of each parameter, and computing the Gelman-Rubin (R-hat) convergence statistic. The data that were fed into the model were less stringently preprocessed than for the saccade latency analysis. Specifically, neither the first saccade was required to be directed to the eventually fixated target, nor were error trials excluded, because the DDM utilizes both correct and incorrect trials to model reaction time distributions.

The best model was selected based on the lowest deviance information criterion (DIC). Even though the DIC penalizes increased numbers of parameters in a model, it still has a bias to prefer more complex models (Wiecki et al., 2013). Therefore, it should only be used as a heuristic for model selection. Condition-specific differences in the parameters of the selected model were analyzed using a Bayesian approach, that is, we sampled from the posterior distributions of the parameters and compared the likelihood of samples being lower in one condition relative to the other. We considered values larger than 97.5% or smaller than 2.5% significant. Note, even though these posterior probabilities are not the same as confidence intervals, they can be interpreted in a similar way (Wiecki et al., 2013). To test the predictive quality of the model, we compared actual data to simulated data, sampled from the posterior distribution of the fitted model and evaluated the correspondence across several summary statistics.

### 2.7 Functional MRI preprocessing

FMRI data was preprocessed using FMRIPrep version 1.0.8 (Esteban et al., 2018), a Nipype (Gorgolewski et al., 2011, 2018) based tool. Each T1w volume was corrected for intensity non-uniformity using N4BiasFieldCorrection v2.1.0 (Tustison et al., 2010) and skull-stripped using antsBrainExtraction.sh v2.1.0 (using the OASIS template). Brain surfaces were reconstructed using recon-all from FreeSurfer v6.0.0 (Dale, Fischl, & Sereno, 1999), and the brain mask estimated previously was refined with a custom variation of the method to reconcile ANTs-derived and FreeSurfer-derived segmentations of the cortical gray-matter of Mindboggle (Klein et al., 2017). Spatial normalization to the ICBM 152 Nonlinear Asymmetrical template version 2009c (Fonov, Evans, McKinstry, Almli, & Collins, 2009) was performed through nonlinear registration with the antsRegistration tool of ANTs v2.1.0 (Avants, Epstein, Grossman, & Gee, 2008), using brain-extracted versions of both T1w volume and template. Segmentation of cerebrospinal fluid (CSF), white-matter (WM) and gray-matter (GM) was performed on the brain-extracted T1w using fast (Y. Zhang, Brady, & Smith, 2001) in FSL v5.0.9 (Jenkinson, Bannister, Brady, & Smith, 2002).

Functional data were motion corrected using mcflirt (FSL v5.0.9). “Fieldmap-less” distortion correction was performed by co-registering the functional image to the same-subject T1w image with inverted intensity (Huntenburg, 2014; Wang et al., 2017) and constrained with an average fieldmap template (Treiber et al., 2016), implemented with antsRegistration (ANTs). This was followed by co-registration to the corresponding T1w using boundary-based registration (Greve & Fischl, 2009) with 9 degrees of freedom, using bbregister (FreeSurfer v6.0.0). Motion correcting transformations, field distortion correcting warp, BOLD-to-T1w transformation and T1w-to-template (MNI) warp were concatenated and applied in a single step using antsApplyTransforms (ANTs v2.1.0) using Lanczos interpolation. After the preprocessing with FMRIPrep, functional data were further high-pass filtered at 1/50 Hz using the fslmaths implementation of Nipype.

Physiological noise regressors were extracted applying CompCor (Behzadi, Restom, Liau, & Liu, 2007). Principal components were estimated for anatomical CompCor (aCompCor). A mask to exclude signal with cortical origin was obtained by eroding the brain mask, ensuring it only contained subcortical structures. For aCompCor, six components were calculated within the intersection of the subcortical mask and the union of CSF and WM masks calculated in T1w space, after their projection to the native space of each functional run. Frame-wise displacement (Power et al., 2014) was calculated for each functional run using the implementation of Nipype. Many internal operations of FMRIPrep use Nilearn (Abraham et al., 2014), principally within the BOLD-processing workflow. See https://fmriprep.readthedocs.io/en/1.0.8/workflows.html for more detail on the pipeline.

### 2.8 Functional MRI analysis

#### 2.8.1 General Linear model

To examine brain activity related to our experimental conditions, we ran an event-related general linear model on the whole brain, separately for each run of each subject. Prior to modeling, functional time series were spatially smoothed with a 5 mm FWHM Gaussian kernel with a nipype implementation of SUSAN (S. M. Smith & Brady, 1997). We separately modeled all combinations of our experimental conditions (both-targets/switch, one-target/switch, both-targets/repeat, one-target/repeat) as well as error trials and the response to the cue display, using display onset times relative to the start of the run as event onset times and the response times as event durations. These events were convolved with a canonical hemodynamic response function (double-gamma), and, together with a number of nuisance regressors, formed the design matrix. Nuisance regressors include the temporal derivative of each event type, motion-related parameters (three regressors each for translation and rotation), framewise displacement (FD), and six anatomical noise regressors (aCompCor). Finally, all volumes with a FD value greater than 0.9 were treated as motion-related outliers and censored, that is effectively excluded from the model. Finally, the data was prewhitened with an autoregressive model to account for temporal autocorrelation (Woolrich, Ripley, Brady, & Smith, 2001). The resulting *t*-statistic maps were combined across runs within participants in a fixed-effect analysis. Next, group analysis was performed with threshold-free cluster enhancement (tfce; S. M. Smith & Nichols, 2009), a voxel-based type statistic that combines the height and spatial extent of local activations. The transformed p-value maps were corrected with nonparametric permutation testing (5000 permutations) as implemented in FSL’s *randomise* (Winkler, Ridgway, Webster, Smith, & Nichols, 2014). Finally, the corrected maps were registered to the FreeSurfer surface (*fsaverage*) coordinate system, using registration fusion (Wu et al., 2018). All reported results were initially thresholded at α = .05, however, for illustration purposes, activity maps were also thresholded at the more stringent α = .01 and shown as overlays (all applied to p-values corrected for multiple comparisons as per above).

#### 2.8.2 Deconvolution

To gain further insight into the time course and extent of the BOLD response as triggered by each event type, we ran a deconvolution analysis in brain regions that are typically considered part of the multiple-demand (MD) network, that has been associated with cognitive control in a variety of contexts (Duncan, 2010; Fedorenko, et al., 2013), plus showed generic switch-related activity in our GLM analysis. To do this, we first converted the preprocessed functional data to represent the percent signal change of the time series, concatenated them across runs and averaged the resulting series within each region of interest (ROI). We defined ROIs based on an existing set of masks of MD subregions (Fedorenko et al., 2013). These ROIs were then combined with voxels that showed significant switch-related activity (collapsed over target availability) in our GLM analysis, yielding 23 ROIs in total. The deconvolution was performed with nideconv (de Hollander & Knapen, 2018). We used the same regressors as in the GLM analysis with the exception that the temporal derivative regressors were not included. All other regressors were convolved with a Fourier basis set, comprising an intercept and four sine-cosine pairs. Prior to estimation, the design matrix was oversampled 20-fold to improve the temporal resolution. For each regressor, beta weights were estimated with ridge regression for each participant separately. The deconvolved time series were extracted from the beta weights and averaged across participants. To statistically test for significant activations, we used a permutation test with 1000 permutations (MNE - one-sample *t*-test; Gramfort et al., 2013) and a cluster-based approach to correct for multiple comparisons. Finally, to test for potential onset differences between proactive and reactive switch-related activity, we used fractional peak latency in combination with the jackknife approach (Liesefeld, 2018, Luck, 2014; J. Miller, Patterson, & Ulrich,1998). In doing so, we averaged the deconvolved time series of all but one participant and identified the time point at which the time series reached 50% of the peak, separately for the proactive and the reactive conditions. To mitigate the influence of local extreme values on the latency estimation, for every time point we averaged the amplitude over five time consecutive points (centered at the current time point). We repeated this procedure leaving out each participant once and computed a paired-sample t-test over the onset estimates. To correct for the artificially reduced error term in the jackknife approach, we followed Miller et al. (1998) by effectively dividing the *t*-statistic by the degrees of freedom.

## 3. Results

### 3.1 Behavioral results

We observed switch costs (longer saccade latency after target switches than target repetitions) only in one-target available blocks but not in both-targets available blocks (Fig. 1B-C). This was statistically confirmed by a two-way repeated-measures ANOVA with target availability (both-targets available vs. one-target available) and transition type (repeat vs. switch) as factors on saccade latency. This ANOVA revealed significant main effects of target availability (*F*(1, 18) = 18.3, *p* < .001, *η^2^* = .50) and transition type (*F*(1, 18) = 23.3, *p* < .001, *η^2^* = .56), and a significant interaction between them (*F*(1, 18) = 16.7, *p* < .001, *η^2^* = .48). A Bayes Factor analysis confirmed this pattern by showing that the model including both main effects and the interaction effect explains the data best (*BF* = 7.8 x 10^5^) and was 12.2 times as likely as the next best model including only the main effects. Overall, saccade latencies were lower in both-targets blocks than in one-target blocks, and lower on switch trials than on repeat trials. Critically however, significant switch costs emerged only in one-target blocks (target repeat: 388 ms vs. target switch: 452 ms; *t*(18) = 5.1, *p* < .001, *Cohen’s d* = 0.63), and not on both-targets blocks (target repeat: 374 ms vs. target switch: 388 ms; *t*(18) = 2.0, *p* = .06, *Cohen’s d* = 0.20). Bayesian t-tests confirmed this conclusion by providing very strong evidence for the presence of switch costs in the one-target condition (*BF_SwitchCosts_* = 374), but no conclusive evidence for either the presence of absence of switch costs in the both-targets condition (*BF_SwitchCosts_* = 1.2).

Next, we analyzed fixation accuracy, that is, the proportion of trials on which participants fixated a target relative to all trials (see Table 1). The data pattern here confirms the saccade latency results with switch costs in one-target blocks and no switch costs in both-targets blocks, precluding an interpretation in terms of a speed-accuracy tradeoff. To test these results, we ran a two-way repeated measures ANOVA with the same factors on accuracy, which also yielded significant main effects of target availability (*F*(1, 18) = 42.6, *p* < .001, *η^2^* = .70) and transition type (*F*(1, 18) = 37.8, *p* < .001, *η^2^* = .68), as well as a significant interaction between them (*F*(1, 18) = 28.6, *p* < .001, *η^2^* = .61). Again, this was supported by a Bayes Factor analysis indicating that the full model (*BF* = 1.8 x 10^10^) is 1,746 times more likely than the model with only main effects (*BF* = 1.8 x 10^7^).

**Table 1.** Percentage correctly fixated targets for all conditions in all three experiments with within-subject 95% confidence intervals (Morey, 2008).

| Target Availability | Target Switch | Target Repeat |
| --- | --- | --- |
| Both-Targets | 96.1 [95.4, 97.0] | 96.9 [96.2, 97.6] |
| One-Target | 87.2 [85.4, 89.0] | 95.3 [94.6, 96.0] |

Finally, to test whether the different display types (target duplicate vs. distractor duplicate) had an influence on the presence or absence of switch costs in each target availability condition, we ran a three-way repeated measures ANOVA with target availability, transition type and display type as factors on saccade latency. However, neither the main effect display type, nor any of the interactions that included that term were significant (*p* > .12). The full ANOVA results can be found in the supplementary material online.

### 3.2 Hierarchical drift diffusion modeling results

Of all models that we ran (see Table S2), the full model with a variable drift rate for target availability and transition type and a variable boundary separation for target availability performed best in explaining the data, as indicated by the lowest DIC (−5.15 × 10^5^). The next best model was the drift-rate-only model with a DIC of −5.13 × 10^5^. We estimated the posterior probability distributions for the condition-specific drift rates and boundary separations and tested for significance directly on the posterior distributions (see Figure 1E). Using the posterior probabilities, we examined how likely it would be for parameter estimates to be greater in one condition compared to another (P[X > Y]). For drift rates, we compared switch to repeat trials in both target availability conditions separately. Drift rates were significantly higher in repeat trials than in switch trials for one-target blocks (switch: *v_mean_* = 2.31, repeat: *v_mea_*_n_ = 3.70, P[switch > repeat] = 0%). In both-targets blocks drift rates were also higher for repeat trials than for switch trials, but the difference was not as large (switch: *v_mean_* = 3.88, repeat: *v_mean_* = 4.22, P[switch > repeat] = 9%). This suggests that participants needed more time to decide whenever selecting a different target than on the previous trial, particularly when only one target was available. Nevertheless, the higher likelihood for drift rates to be larger for repeat than switch trials when both targets were present suggests that also in this condition, some switch-related cost was present. Even though the best model included separate estimates for boundary separation for one-target blocks and both-targets blocks, comparing these boundary separation estimates to each other yielded virtually the same value (one-target available: *a_mean_* = 1.17, both-targets available: *a_mean_* = 1.16, P[both-targets > one-target] = 47%). This suggests that participants did not adjust their response caution across target availability conditions. This finding is somewhat unintuitive given that the best model included a separate boundary separation parameter for each target availability condition, but could be caused by the DIC being biased toward the more complex model (see Wiecki, 2013). Finally, to confirm that the full model accurately captured the data, we examined the quality of the model fit. In addition, we also checked whether the simpler drift-rate-only model (only drift rate could vary across experimental conditions) was also representative of the data, we analyzed that model as well. For this purpose, we generated data by sampling from the posterior distributions of the parameters and compared the simulated data to the original data. Importantly, key summary statistics, such as accuracy, mean, median, quantiles (10, 30, 50, 70, 90) saccade latencies, were all recovered in the simulated data, as indicated by all summary statistics lying within the 95% credible intervals. This indicates that both models provide a good fit to the data and interpreting their parameters is warranted.

### 3.3 Neuroimaging results

#### 3.3.1 General linear model

To determine whether the behavioral effects in terms of switch costs are governed by separable cognitive control mechanisms, we used a general linear model (GLM) to examine whether BOLD activity associated with updating a target representation depended on how many unique targets were available in a search display. Before comparing switch-related activity across target availability conditions, we first contrasted switch trials with repeat trials, separately within the one-target and the both-targets condition. When both targets were available, switches elicited widespread activations across cerebral cortex and cerebellum, in a network reminiscent of the multiple-demand (MD) network (see Fig. 2 and Table 2). Bilateral frontal activations include dorsolateral prefrontal cortex (dlPFC), frontopolar cortex, dorsal premotor cortex (PMd), inferior frontal junction (IFJ), anterior insula / frontal operculum cortex (aINS), posterior cingulate gyrus (pCG), and medial frontal cortex (spanning from the dorsal anterior cingulate cortex to the frontal eye fields, mFC/dACC). Parietal activations were found bilaterally in the superior parietal lobule (SPL), extending across the intraparietal sulcus (IPS) into the inferior parietal lobule (IPL), including supramarginal and angular gyrus. In the occipital and temporal lobe, there was switch-related activity in the intracalcarine sulcus and in temporo-occipital regions, bilaterally and in the right inferior temporal gyrus. Finally, several subregions in the cerebellum were activated as well. When only one target was available, fewer significant clusters were found (see Fig. 3 and Table 2). Activations were restricted primarily to posterior regions in the parietal and occipital lobe, including IPS, SPL, and IPL as well as the cerebellum. However, two smaller activated regions were also found in the left IFJ, and the dlPFC at the border with the frontopolar cortex.

**Figure 2.**
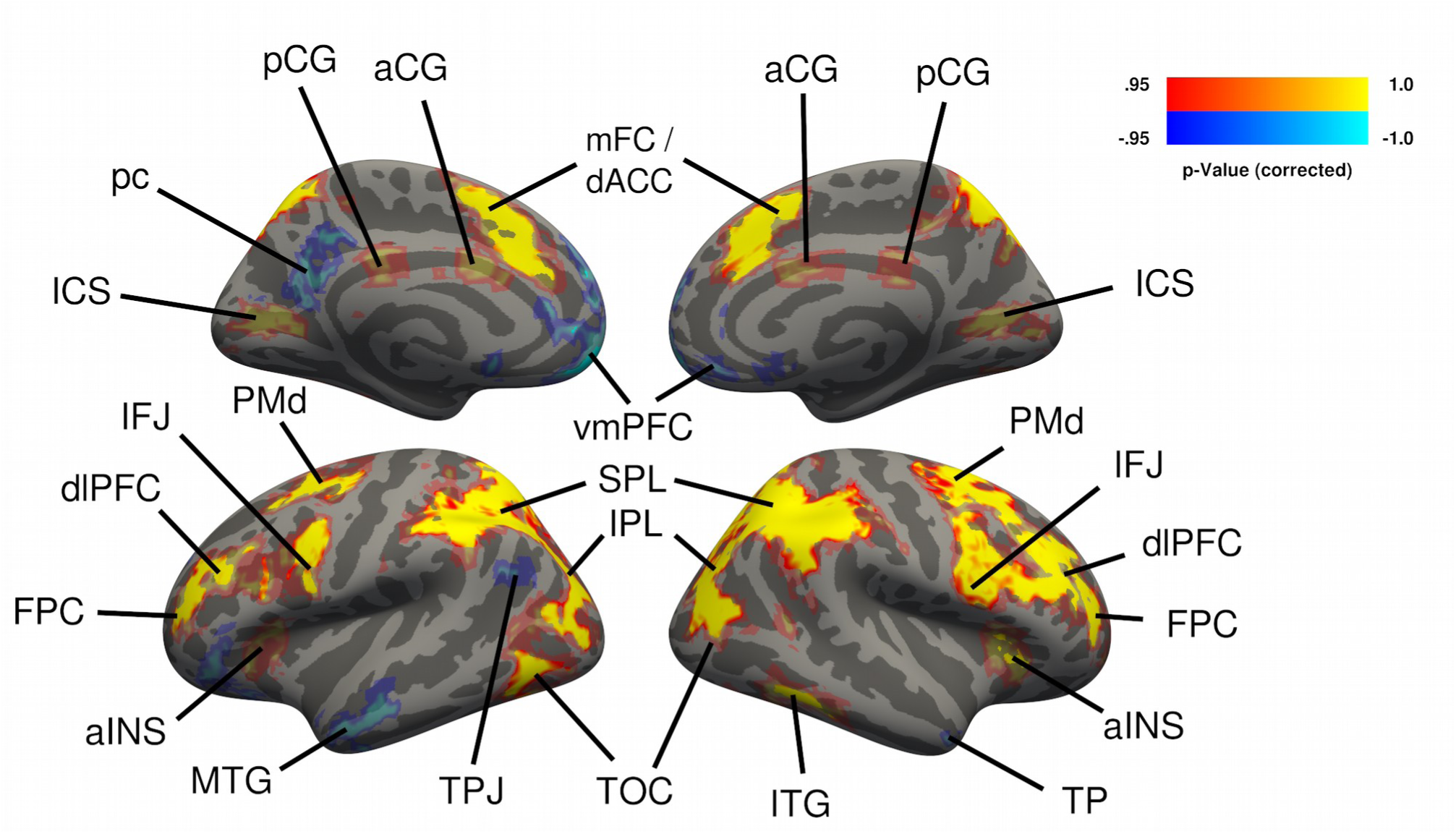
Cerebral activations for the free choice condition (proactive control demand). Activations shown in yellow-red represent the contrast free switch > free repeat; activations shown in blue represent the contrast free repeat > free switch. Group-level *t*-statistics maps were computed with the tfce-method (S. M. Smith & Nichols, 2009) and corrected for multiple comparisons using nonparametric permutation testing. The resulting P-value maps were thresholded at α = .05 and projected onto the fsaverage surface using registration fusion (Wu et al., 2018), with translucent coloring. In addition, regions that were also significant at α = .01 are shown in saturated colors. Free switches were associated with higher activity than free repeats across both hemispheres in dorsolateral prefrontal cortex (dlPFC), frontopolar cortex (FPC), dorsal premotor cortex (PMd), inferior frontal junction (IFJ), anterior insula/frontal operculum cortex (aINS), posterior cingulate gyrus (pCG), anterior cingulate gyrus (aCG), medial frontal cortex (spanning from the dorsal anterior cingulate cortex to the frontal eye fields, mFC/dACC), superior parietal lobule (SPL), inferior parietal lobule (IPL), intracalcarine sulcus (ICS), right inferior temporal gyrus (ITG), temporo-occipital cortex (TOC), and bilateral cerebellum (not shown here). Free repeats were associated with higher activity than free repeats in the left precuneus (pc), bilateral ventromedial prefrontal cortex (vmPFC), left medial temporal gyrus (MTG), left temporoparietal junction (TPJ) and right temporal pole (TP).

**Figure 3.**
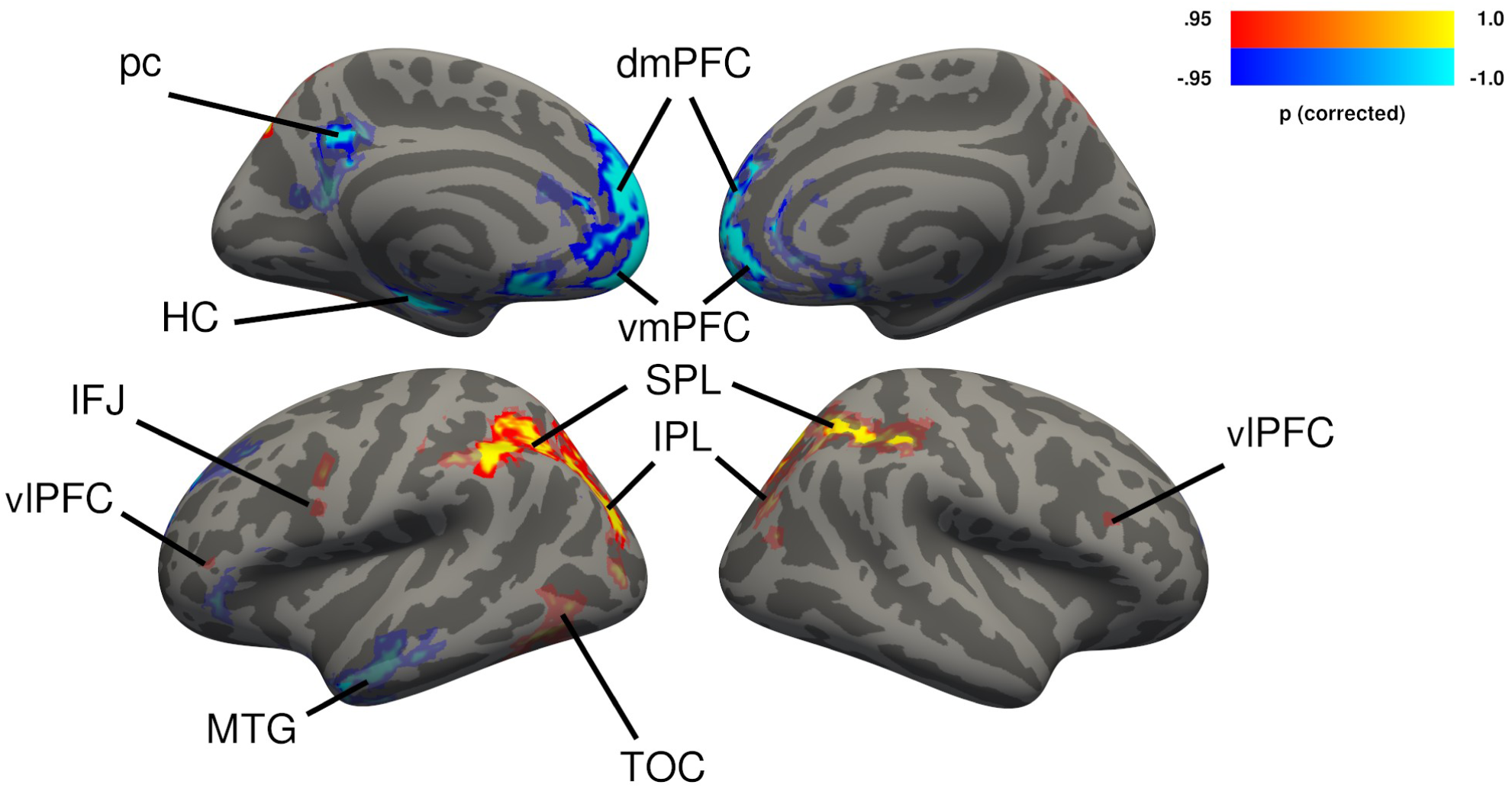
Cerebral activations for the imposed choice condition (reactive control demand). Activations shown in yellow-red represent the contrast imposed switch > imposed repeat; activations shown in blue represent the contrast imposed repeat > imposed switch. Group-level *t*-statistics maps were computed with the tfce-method (S. M. Smith & Nichols, 2009) and corrected for multiple comparisons using nonparametric permutation testing. The resulting P-value maps were thresholded at α = .05 and projected onto the fsaverage surface using registration fusion (Wu et al., 2018), with translucent coloring. In addition, regions that were also significant at α = .01 are shown in saturated colors. Imposed switches were associated with higher activity than imposed repeats bilaterally in dorsolateral prefrontal cortex (dlPFC), left inferior frontal junction (IFJ), superior parietal lobule (SPL), inferior parietal lobule (IPL), temporo-occipital cortex (TOC), and bilateral cerebellum (not shown here). Imposed repeats were associated with higher activity than imposed repeats in the left precuneus (pc), bilateral ventromedial (vmPFC), dorsomedial prefrontal cortex (dmPFC), left medial temporal gyrus (MTG), and left hippocampus (HC).

**Table 2.**
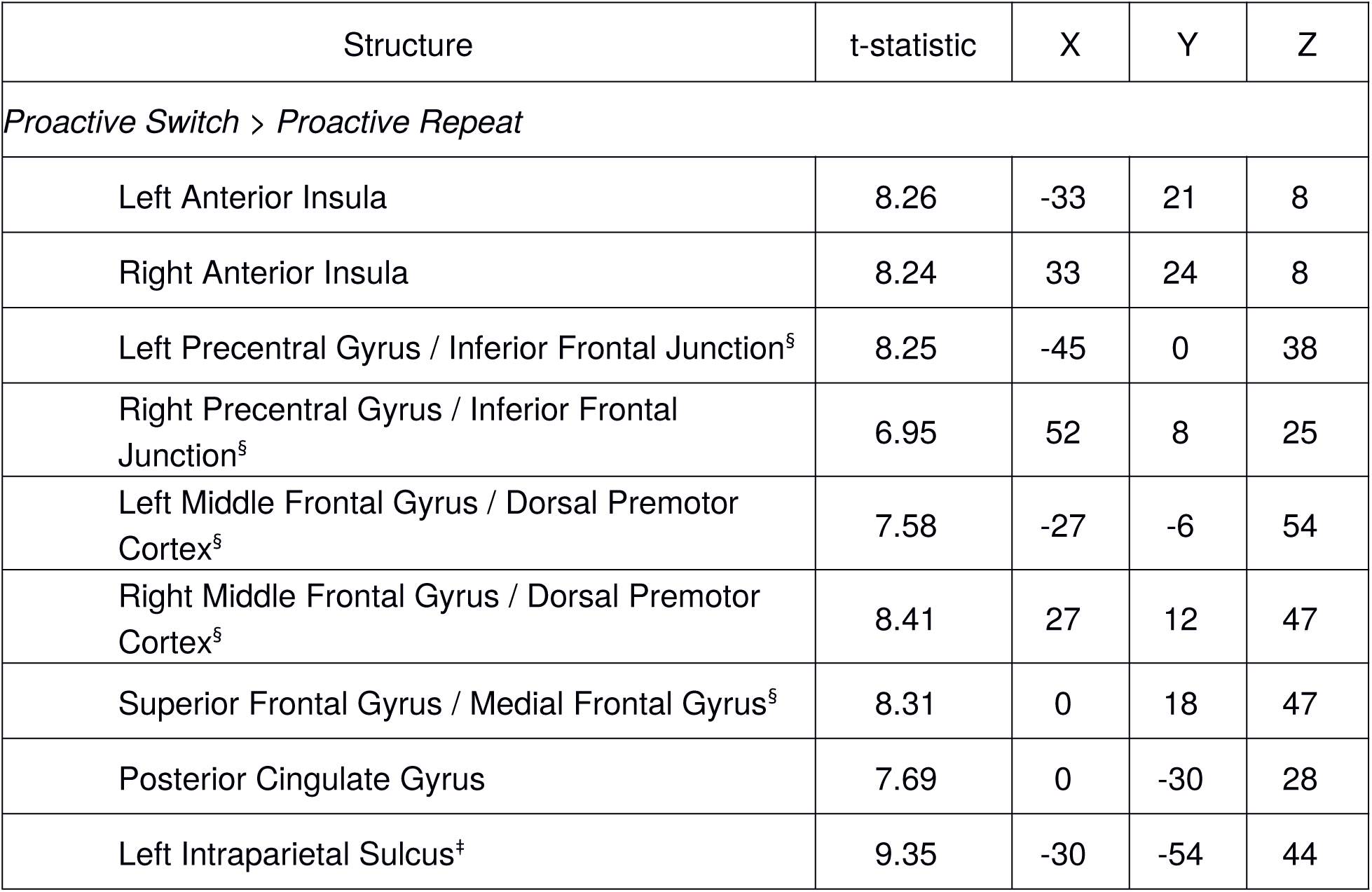
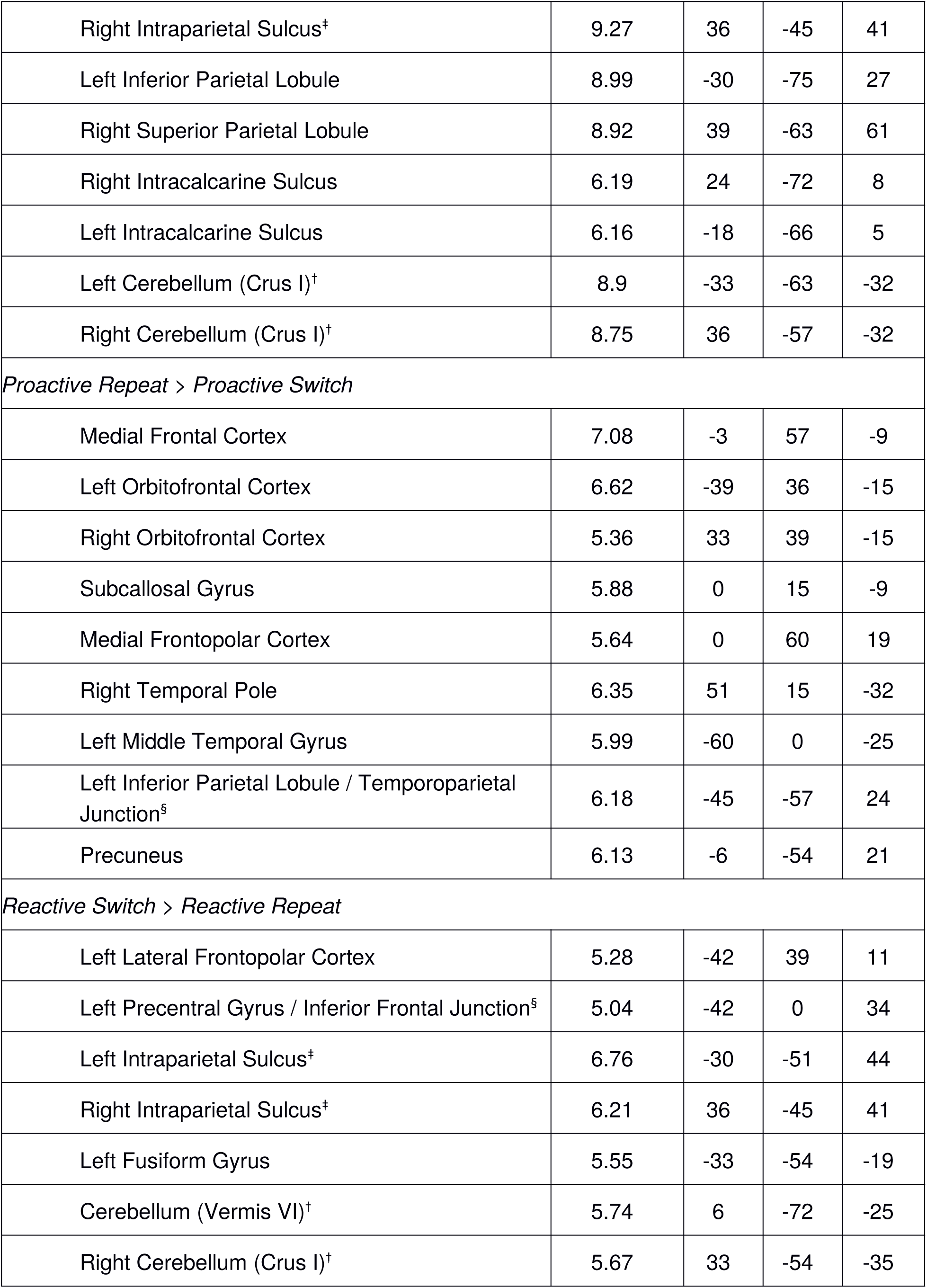
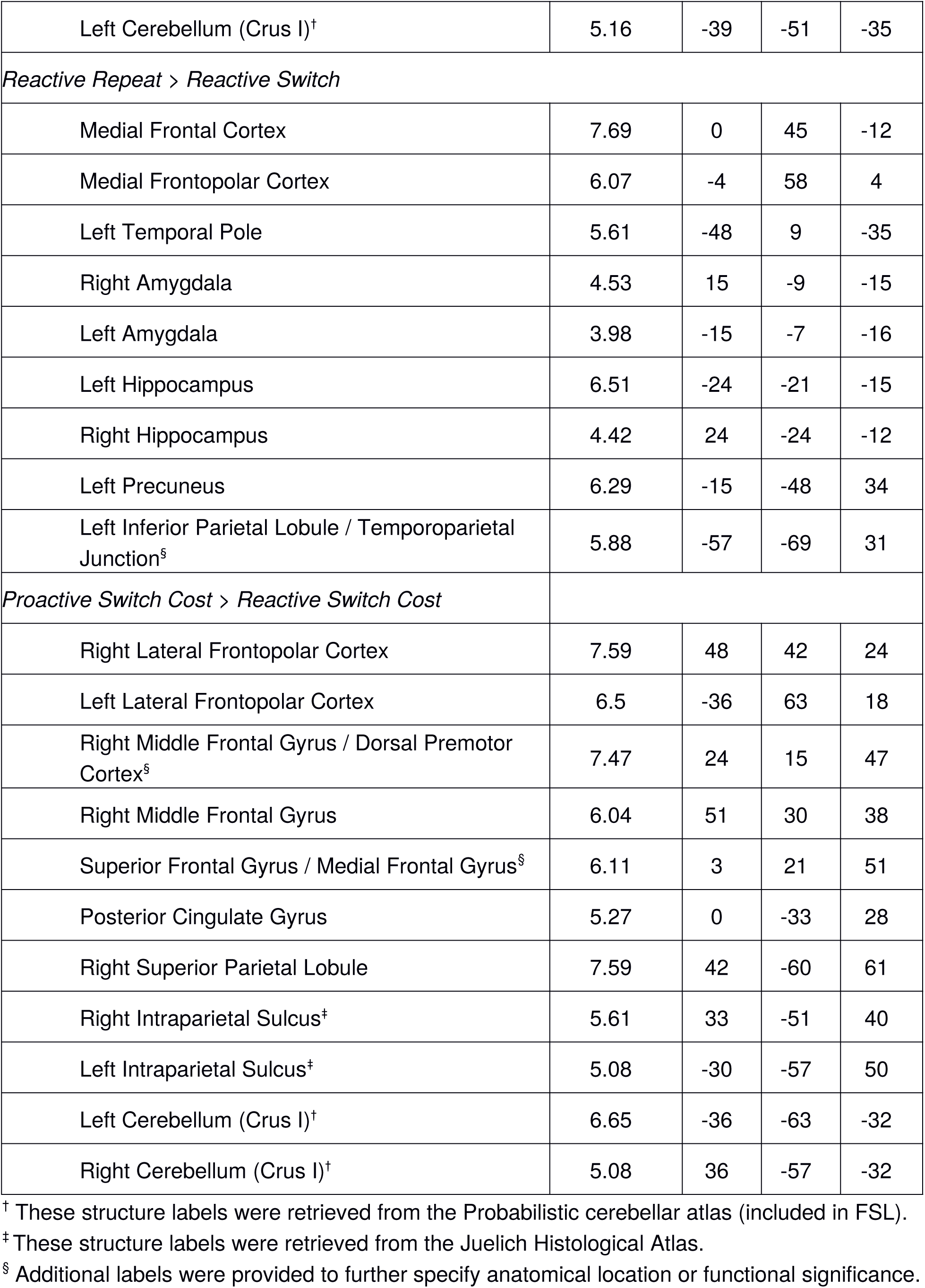
Localization of activations for the main contrasts. Coordinates of local maxima are reported in MNI152-space. Large clusters were split into subclusters based on anatomical considerations. Structure labels are based on the Harvard-Oxford anatomical atlas.

Next, to statistically compare switch-related activity between target availability conditions, we directly compared activity associated with each target availability condition to each other. However, because target availability was manipulated at the block-level, when directly comparing switch-related activity across target availability conditions, we might pick up on overall block differences rather than true switch-related differences. We therefore computed a double contrast, in which we first isolated switch-related activity per target availability condition by subtracting repeat activity from switch activity, and next, contrasted these differences to obtain the neural correlate of switching when both targets were available versus when one target was available. When considering regions where switch-related activity was stronger in both-targets blocks than in one-target blocks, we again found activations closely resembling the MD network, including bilateral dlPFC, frontopolar cortex, mFC/dACC, pCG, SPL, IPS, and Cerebellum (Fig. 4). The opposite contrast–more switch-related activity in the one-target condition than in both-targets available condition–yielded no significant activations. One possibility might be that in the one-target condition, target representations often needed to be updated both on switch and repeat trials, for example if participants did not anticipate any of the targets. If this is the case, switch and repeat trials would be more similar in the one-target available condition, so that activity reflecting switch costs would be reduced.

**Figure 4.**
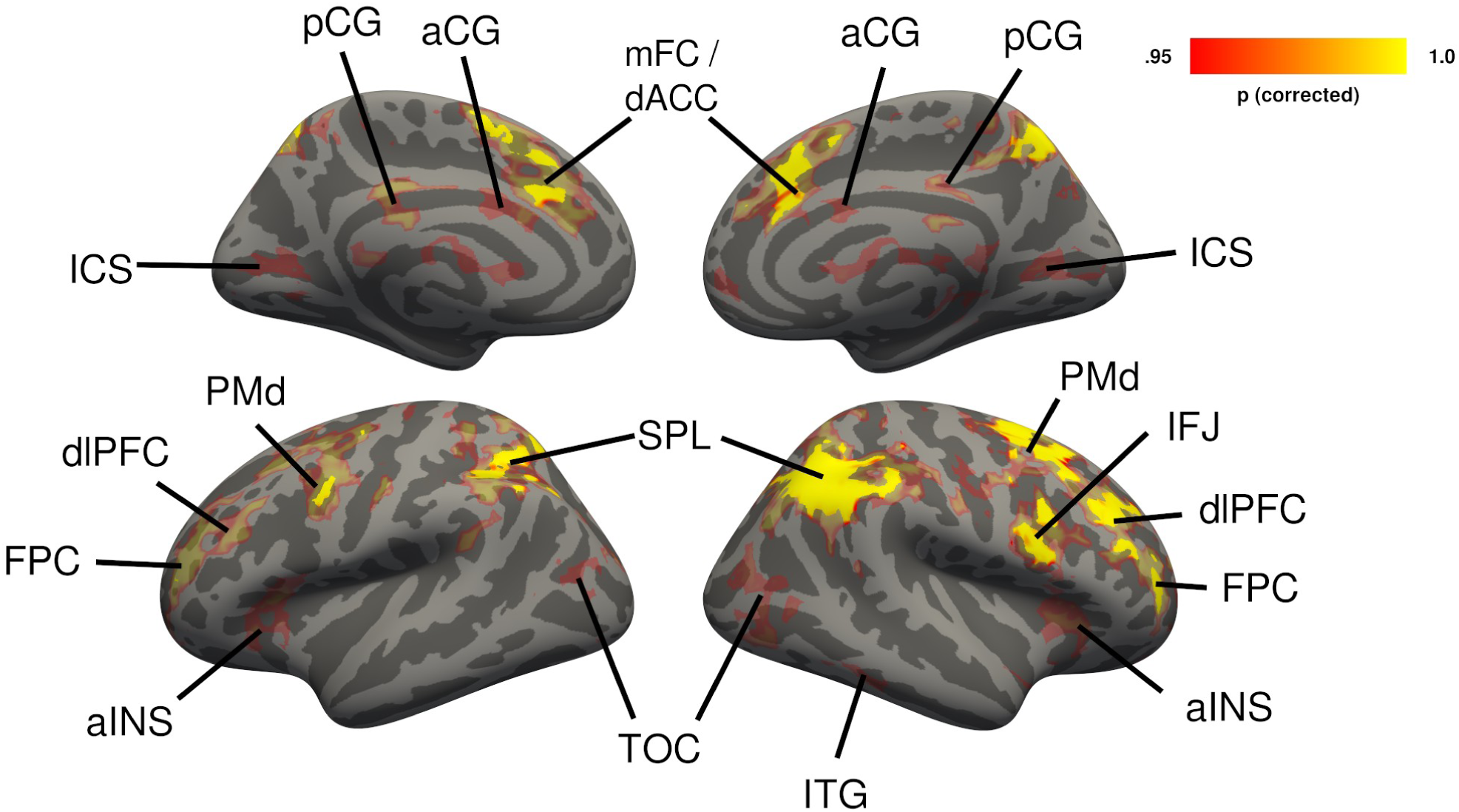
Cerebral regions in which free switch cost (free switch > free repeat) yielded stronger activity than imposed switch cost (imposed switch > imposed repeat), shown in yellow-red. Group-level *t*-statistics maps were computed with the tfce-method (S. M. Smith & Nichols, 2009) and corrected for multiple comparisons using nonparametric permutation testing. The resulting P-value maps were thresholded at α = .05 and projected onto the fsaverage surface using registration fusion (Wu et al., 2018), with translucent coloring. In addition, regions that were also significant at α = .01 are shown in saturated colors. Free switch costs were associated with higher activity than imposed switch cost across both hemispheres in dorsolateral prefrontal cortex (dlPFC), frontopolar cortex (FPC), dorsal premotor cortex (PMd), inferior frontal junction (IFJ), anterior insula/frontal operculum cortex (aINS), bilateral posterior cingulate gyrus (pCG), anterior cingulate gyrus (aCG), medial frontal cortex (mFC/dACC), superior parietal lobule (SPL), inferior parietal sulcus (IPS), intracalcarine sulcus (ICS), right inferior temporal gyrus (ITG), temporo-occipital cortex (TOC), and bilateral Cerebellum (not shown here).

To investigate this possibility, we directly compared switch activations between target availability conditions without taking repeat events into account (see Fig. S2). For the contrast both-targets switch greater than one-target switch, a very similar pattern was found as in the double-contrast analysis (only bilateral aINS and Caudate were additionally active), thus confirming the previous findings and suggesting that block differences do not seem to play an important role. More importantly, when considering the opposite contrast, significant clusters of activation were now found in the left ventrolateral prefrontal cortex (vlPFC), precuneus, and left temporoparietal junction, regions that have been considered part of the default mode network (DMN; Raichle, 2015; Raichle et al., 2001). The DMN has recently been linked to automated behavior, not rigorously governed by cognitive control (Vatansever, Menon, & Stamatakis, 2017). Therefore, stronger DMN activations in one-target blocks could be explained by less control being applied during switches in this condition compared to both-targets blocks. If so, the same should be true for repeat trials. For this reason, these DMN activations might not have emerged in the double-contrast procedure, as these activations canceled each other out when switch and repeat trials in the one-target condition were contrasted with each other. To investigate whether there actually was DMN activity associated with repeat trials, in an exploratory analysis, we examined whether there were regions in which repeat events led to stronger activation than switch events, separately for the both-targets and one-target conditions.

In this exploratory analysis, we effectively reversed the contrast that was used to isolate switch-related activity in the first step of the double contrast procedure. We indeed detected strong activity along the medial wall of the PFC, the orbitofrontal cortex (OFC), the precuneus, the left medial temporal gyrus (MTG), and the temporoparietal junction (TPJ) across both target availability conditions (opposite contrast shown in blue in Fig. 2 and Fig. 3). In addition to these common activations, the amygdala and the hippocampus were selectively active in the one-target blocks (Fig. 3). Directly comparing both-targets to one-target repeat trials (not taking switch trials into account), yielded three cluster in which there was stronger activity in the one-target condition. These clusters were located in the precuneus, the medial PFC, MTG, and the TPJ (Fig. S3). The opposite contrast did not show any significant activations, suggesting that the DMN was activated more strongly when one target was available than when both targets were available, which could indicate a higher demand for cognitive control during both-targets blocks than during one-target blocks (see Discussion), or conversely, more automated behavior during the latter. To test this hypothesis, finally, we compared activity between target availability conditions across all event types (collapsed across transition type). This analysis indicates primarily DMN activity (precuneus, vlPFC, TPJ, and medial PFC) in one-target blocks and MD network activity (bilateral dlPFC, aINS, PMd, IFJ, frontopolar cortex, mFC/dACC, SPL) in both-targets blocks (Fig. S4), in line with the hypothesis that more control is demanded during both-targets blocks.

Taken together, the GLM findings demonstrate that for both-targets as well as one-target switches, activations were found in what is known as the multiple-demand network. However, these activations were stronger and more widespread for free switches than for imposed switches. Furthermore, during repeat trials, the DMN was strongly active, particularly during one-target blocks. Irrespective of transition type (switch versus repeat), the MD network seems to be more engaged when both targets are available than when only one is there, in which case the DMN is predominantly active.

#### 3.3.2 Deconvolution analysis

The standard approach of modeling the BOLD response with a canonical hemodynamic response function (HRF) maximizes sensitivity for activations at the expense of being more biased towards a predefined shape of the response (Poldrack, Mumford, & Nichols, 2011). To characterize potential interregional variability and accommodate non-standard BOLD-responses not captured by the double-gamma function that we used in the GLM approach, we employed a deconvolution analysis. Deconvolution has the advantage that the shape and the time course of the HRF can vary and some temporal information can be retained. We limited this analysis to ROIs that are part of the MD network and showed switch-related activity (collapsed across target availability) in the GLM (see section 2.8.2), to limit the number of analyses, while still considering most regions in which switch-related activity might be found. Across both cerebral cortex and cerebellum, 23 ROIs were considered. We focused on activation differences between imposed and free switches, specifically where the activation patterns diverge from the standard GLM results described above. Such differences were found primarily for imposed switches. Specifically, the deconvolution identified significant activity bilaterally in the anterior Insula (aINS) and the left dorsal premotor cortex (PMd) located on the superior frontal gyrus. Nevertheless, in these regions, free switches still elicited a stronger response than imposed switches (see Fig. 5). Across all other ROIs, we observed four patterns: regions with neither imposed nor free switch activity, regions with only free switch activity, regions with both imposed and free switch activity but more activity on free switches, and regions with equal amounts of imposed and free switch activity. However, in all those regions the deconvolution yielded the same qualitative pattern as the standard GLM approach.

**Figure 5.**
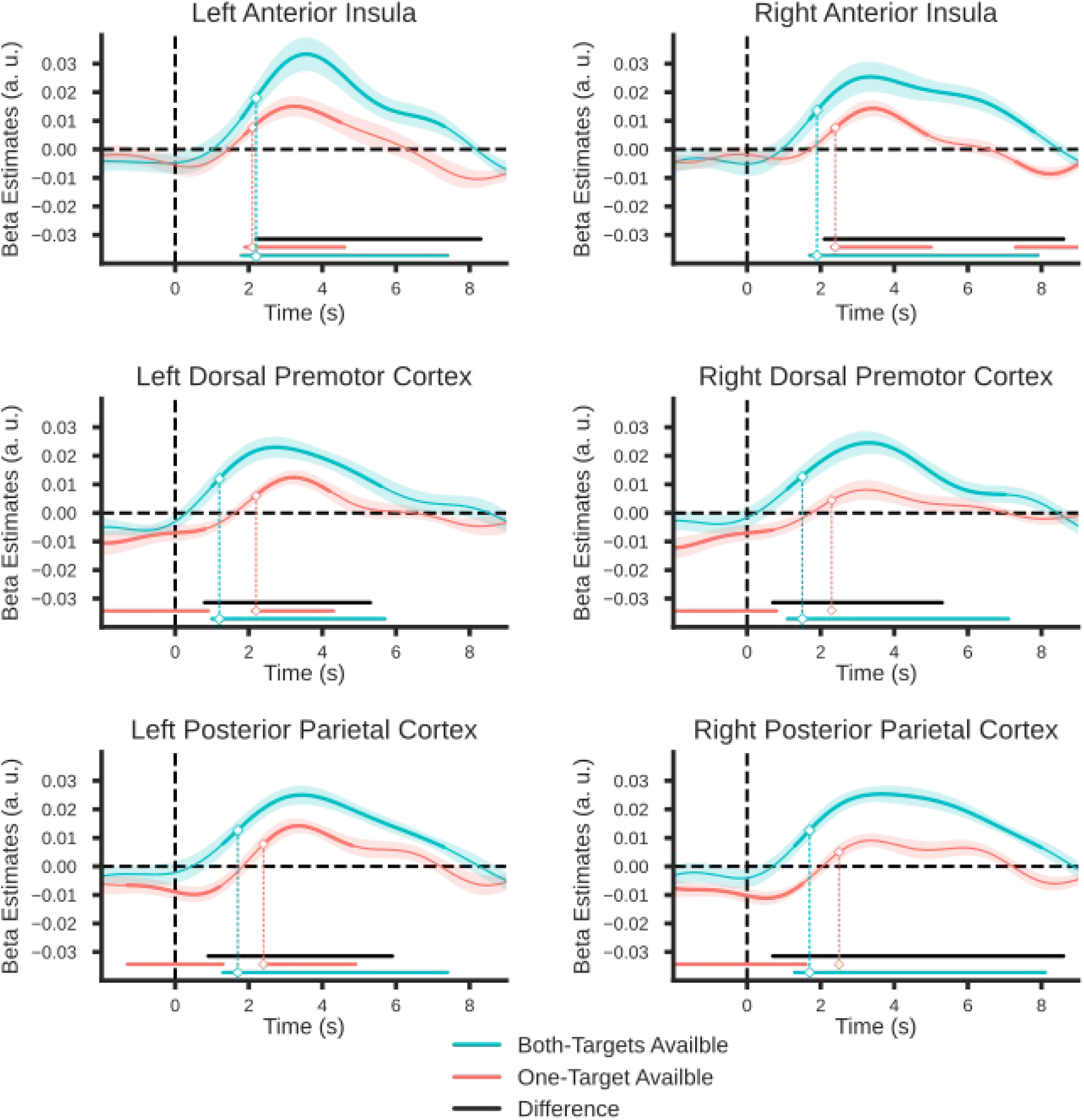
Group-averaged beta estimates of neural activation time course in selected regions of interest. Deconvolution analysis was used to model the BOLD response for each event type separately. For each target availability condition, the difference of the time courses for switch and repeat trials was computed and the resulting time courses are shown here. The shaded color bands represent 68% confidence intervals (±1 SEM). Thick lines as well as horizontal bars indicate significant clusters (at α = .05) as produced by cluster-based permutation testing (5000 permutations). The black horizontal bars indicate the range over which the difference between the target availability condition was significant. The marked time points (vertical dashed lines) indicate the latency of 50% maximum amplitude as estimated using a jackknife approach, as a measure of the onset of activation (Miller, Patterson and Ulrich, 1998; Luck, 2014; Liesefeld, 2018).

Note that beyond regional differences between imposed and free switches, these two conditions could also differ in temporal aspects. In fact, a strong prediction of the dual mode of control framework is that proactive control should begin before trial onset, whereas reactive control should only be invoked after the search display onset. Van Driel and colleagues (2019) provided strong support for this prediction using a very similar paradigm to ours in combination with more time-sensitive EEG measures. To test for potential onset differences also in the fMRI signal, we measured the estimated onset latency in combination with a jackknife approach. The results yielded significantly earlier proactive switch activity in the left PMd (M_proactive_ = 1210 ms, M_reactive_ = 2237 ms, *t_c_*(18) = 2.77, *p_c_* = .01), right PMd (M_proactive_ = 1484 ms, M_reactive_ = 2300 ms, *t_c_*(18) = 2.53, *p_c_* = .02), left IFJ (M_proactive_ = 1484 ms, M_reactive_ = 2226 ms, *t_c_*(18) = 2.79, *p_c_* = .01), right IFJ (M_proactive_ = 1505 ms, M_reactive_ = 2000 ms, *t_c_*(18) = 2.99, *p_c_* = .008), left posterior parietal cortex (M_proactive_ = 1721 ms, M_reactive_ = 2405 ms, *t_c_*(18) = 2.22, *p_c_* = .04), right posterior parietal cortex (M_proactive_ = 1747 ms, M_reactive_ = 2516 ms, *t_c_*(18) = 3.33, *p_c_* = .004), and mFC/dACC (M_proactive_ = 1284 ms, M_reactive_ = 2026 ms, *t_c_*(18) = 2.79, *p_c_* = .01), but no such difference in the bilateral aINS (left: M_proactive_ = 2152 ms, M_reactive_ = 2115 ms, *t_c_*(18) = 0.41, *p_c_* = .69; right: M_proactive_ = 1900 ms, M_reactive_ = 2410 ms, *t_c_*(18) = 1.50, *p_c_* = .15). Some of these areas, notably mFC/dACC and PMd, are consistent with the midfrontal topography of the free choice related beta-oscillatory suppression that was observed by van Driel et al (2019). Note that one might also expect to find onset differences in the frontopolar cortex, given its presumed role in voluntary switching (Mansouri et al., 2017, Pollmann, 2016). However, as there was virtually no activity related to imposed switches in this region, onset difference could not meaningfully be determined.

## 4. Discussion

In this study we set out to examine which brain regions are recruited in multiple-target search, depending on whether observers are free to select a target or whether they are forced to select a particular target. For this purpose, we asked observers to look for multiple targets and we manipulated whether both or only one of the two potential target colors were present in a search display. We reasoned that the presence of both targets would enable observers to use proactive control to prepare a search, whereas the presence of only a single item would require reactive control whenever the observer expected the wrong target. In accordance with previous findings (Ort et al., 2017, 2018), we found clear switch costs in terms of both saccade latency and saccade accuracy when only one target category was present in a search display, while there were no switch costs when both targets were available. This finding is further supported by the results of hierarchical drift diffusion modeling, which revealed lower drift rates on switch compared to repeat trials when one target was available. When both targets were present, drift rates were also lower for switch than for repeat trials, but this difference was much smaller than in the one-target condition. This suggests that observers used the predictability of the both-targets condition to prepare selection of either one of the targets, so that potential costs associated with updating the currently active target representations remained latent.

Importantly, using fMRI measures we provide new evidence regarding the neural mechanisms underlying these switches of feature-based attention. We found the frontoparietal multiple-demand network (Duncan, 2010; Fedorenko, et al., 2013) to be strongly associated with free target switches. Imposed target switches elicited a similar, yet weaker activity pattern in the posterior parietal cortex (PPC), and relatively smaller activity clusters in frontal regions at the inferior frontal junction (IFJ) and dorsolateral prefrontal cortex (dlPFC). Furthermore, the direct comparison of free and imposed switches indicates that the multiple-demand network is more strongly involved during free than during imposed switches. In contrast, parts of the default mode network are activated stronger in blocks involving imposed switches. Assuming that target availability conditions primarily differed with respect to whether observers used proactive control (both-targets available) or reactive control (one-target available), our findings suggest that these two modes of control can indeed be dissociated during multiple-target search. More specifically, by means of a deconvolution analysis, we were able to categorize these differential activations into regions that exclusively activate for free switches (dlPFC, frontopolar cortex, and medial frontal cortex/mFC) and regions that are also active during imposed switches but to a lesser extent (IFJ, dorsal premotor cortex/PMd, and PPC). Furthermore, these regions activated earlier for free switches, corroborating their role in preparatory cognitive control in anticipation of a demanding event.

The observed activations for imposed switches are reminiscent of earlier reports on stimulus or task-induced feature-based attention shifts with activations primarily located bilaterally in PPC and PMd (Greenberg et al., 2010; Liu et al., 2003; Pollmann et al., 2006, 2000, Slagter et al., 2006, 2007). This also matches the observation that IFJ and PPC are involved in updating and representing task sets across a variety of tasks (Brass & von Cramon, 2004; Kim et al., 2012). In particular, it has been suggested that while IFJ is responsible for updating task-specific information, the PPC maintains such information and implements task sets (Brass & von Cramon, 2004; Bunge, Kahn, Wallis, Miller, & Wagner, 2003; Greenberg et al., 2010; Shulman, 2002; Slagter et al., 2007). Finally, the activity in the anterior insula that we observed after deconvolution analyses of both types of switches may be part of a network that signals salient events (such as the absence of an expected target color) and the need to initiate a cascade of control signals that eventually update the active target representation (Menon & Uddin, 2010; Power & Petersen, 2013; Seeley et al., 2007). We isolated the neural response to feature-based attention shifts from the additional types of changes that may contribute to task-switch costs (e.g. Meiran, 2010), in particular shifts of stimulus-response mapping. Our findings suggest that establishing a new attentional set is a rather “cheap” process that requires only minimal frontal activity (see Ort, Fahrenfort, ten Cate, & Olivers, 2019; Moore & Weissman, 2010 for behavioral and electrophysiological evidence). Furthermore, similar to Gmeindl et al. (2016), we directly compared endogenous, self-initiated target switches to imposed switches, but of feature-based rather than spatial attention. Importantly, our design allowed us to link either switch-related activity to specific events in the experiment. In doing so, we show that there is common, but also distinct neural activity underlying these types of switches.

However, some findings were unexpected, in particular with respect to imposed switches. First, with the standard GLM approach, we did not observe any significant imposed-switch-related activity in the dorsal premotor cortex (PMd; presumably the location of the human frontal eye fields), an area that has previously been shown to be related specifically to feature-based attention shifts (Kim et al., 2012). Using deconvolution, we were able to detect significant, but relatively weak activity in the left PMd. A possible explanation may be that in contrast to earlier studies of feature-based attention shifts (e.g. reviewed in Kim et al., 2012), in our study, there were no changes in the stimulus-response mapping associated with imposed switches. Note that an imposed target shift did not systematically signal a particular eye movement as target location and target identity were unrelated. Therefore, there was no need to activate or update a certain stimulus-response mapping, which has been suggested to be a function of the PMd (e.g. Badre & D’Esposito, 2009; Hopfinger, Buonocore, & Mangun, 2000; Kim et al., 2012). This is also supported by Pollmann and colleagues (2006), who, during a visual search task, separated attention shifts from response shifts and found only the latter to activate the PMd.

Second, unlike Jiang and colleagues (2018), we did not observe any activity in the anterior cingulate cortex (ACC) related to imposed switches. This region has been linked to conflict monitoring in numerous studies (e.g. Botvinick, Nystrom, Fissell, Carter, & Cohen, 1999; Ito, Stuphorn, Brown, & Schall, 2003; Jiang et al., 2015; Kerns et al., 2004; Ullsperger, Danielmeier, & Jocham, 2014). As observers could not anticipate imposed target switches, we also expected a degree of surprise whenever the target changed. This signal has been suggested to be related to conflict-processing and to originate in the medial frontal cortex (e.g. Cavanagh & Frank, 2014). However, experienced conflict in Jiang et al. (2018) may have been stronger due to the fact that participants were explicitly cued as to which task to expect, while in our study any build-up of expectations was left to the observer. Furthermore, unlike in the Jiang et al. study, observers did not have to manage a target-specific stimulus-response mapping in our study. Overall, we believe that target selection in our paradigm was relatively easy and therefore did not evoke strong conflict-related signals in the frontal cortex. That said, in a recent EEG study with a very similar paradigm (van Driel et al., 2019), we did observe a power enhancement in the delta/theta-frequency band after imposed switches over midfrontal electrodes. This signal has been suggested to be related to conflict-processing and to originate in the medial frontal cortex (e.g. Cavanagh & Frank, 2014). It remains to be investigated why we found no corresponding source here.

Some support for the hypothesis that relatively little control was exerted in the imposed target condition comes from the default mode network activity that we observed, particularly for repeat trials. The default mode networkhas recently been shown to not just reflect an idle brain state, but to also activate during various tasks (Elton & Gao, 2015; Konishi, McLaren, Engen, & Smallwood, 2015; Smallwood et al., 2013; V. Smith, Mitchell, & Duncan, 2018; Spreng, 2012; Spreng et al., 2014; Vatansever et al., 2017). Even though its functional significance is still debated, there is increasing evidence that suggests the default mode network is related to internally-generated thought (Konishi et al., 2015), decoupled from immediate sensory input or context-representation (V. Smith et al., 2018). Maybe most importantly, Vatansever and colleagues (2017) demonstrated that even though the cognitive control network is strongly involved in acquiring task rules, once those rules have been learned, the default mode network becomes active while applying them. They concluded that whenever the current task context is predictable, individuals enter a form of “autopilot” mode in which correct responses can be made without explicit cognitive control. We argue the same may happen in our paradigm: During phases of target repeats, participants were able to select the correct target disk in a low-control, automated manner.

The activation patterns associated with free switches are similar to previously reported activations related to proactive control demand (Irlbacher et al., 2014; Jiang et al., 2018). In addition to parietal and posterior frontal activity as was also observed for imposed switches, two key activations are of primary importance here. First, there were strong medial activations spanning from dorsal anterior cingulate cortex (dACC) to the supplementary motor area (SMA). Activity in these regions has been associated with self-generated choice (Demanet et al., 2013; Forstmann et al., 2006; Gmeindl et al., 2016; Orr & Banich, 2013; Passingham et al., 2010; Soon et al., 2008; Taylor et al., 2008; Wisniewski et al., 2016; Wisniewski et al., 2015; J. Zhang et al., 2013). The present pattern of activations matches those findings, consistent with the idea that participants used the available information to prepare a switch trial in advance. Second, activity was found in the lateral frontopolar cortex. This region has been associated with making free decisions, but also with the evaluation of alternative goals in the context of exploratory behavior (Mansouri et al., 2017; Pollmann, 2016). We believe that this activity might reflect participants evaluating whether or not to switch to the other target color during a streak of target repeats. Nevertheless, even though these activations were specific to the free choice condition, they may only indirectly relate to proactive control, inasmuch as this information can be used by a cognitive control system to signal when (and supposedly how much) proactive control should be invoked.

In addition, we also found activity along the dlPFC. In line with an interpretation in terms of proactive control, this activity might reflect preparatory updating and maintaining of task rules (Braver, Paxton, Locke, & Barch, 2009). However, as the dlPFC has been linked to a wide range of executive functions, such as working memory, planning, and inhibition (e.g. Niendam et al., 2012), we cannot exclude that other factors caused the activations in this region. For example, dlPFC activity could have been caused by additional mental effort and working memory demand associated with overall planning or keeping track of the switches, in order to adhere to the task instructions (e.g. Braver et al., 1997; Bunge, Ochsner, Desmond, Glover, & Gabrieli, 2001; Dosenbach et al., 2008; Rypma & D’Esposito, 1999; Shenhav, Botvinick, & Cohen, 2013). Nevertheless, it could be argued such additional cognitive processes, despite not being cognitive control in a strict sentence, are essential for proactive control. In this sense, proactive control requires the maintenance of the current and the targeted state (working memory), planning target selection on future trials (planning) and making the decision to invoke proactive control at a given moment (intention). Therefore, the actual usage of proactive control can be seen as a consequence of an cascade of other cognitive processes.

Beyond that, the present findings provide further support for multiple-state models of working memory postulating that the number of memory items that can concurrently affect behavior at any given moment is limited (Huang & Pashler, 2007; Oberauer, 2002; Olivers, Peters, Houtkamp, & Roelfsema, 2011). The fact that we observed switch costs indicates that observers did not distribute resources equally across multiple target representations. This interpretation is supported by the fMRI results which show switch-related activity in both target availability conditions in regions that have previously been associated with updating of attentional sets (Greenberg et al., 2010; Liu et al., 2003; Pollmann et al., 2006, 2000, Slagter et al., 2006, 2007; Wager et al., 2004), including bilateral posterior parietal cortex and inferior frontal junction. Switch-related activity in these regions suggests that in both target availability conditions switch trials were associated with priority shifts, therefore supporting dynamic weighing of attentional relevance between target representation (see also van Driel et al., 2019).

## 5. Conclusion

We investigated the contributions of proactive and reactive control to target selection during multiple-target search. We found that both control mechanisms activate a similar network that has previously been associated with shifts of feature-based attention, with proactive control eliciting greater activity generally. In addition, proactive switching also activated other frontal regions that have been linked to free choice and evaluating alternative options other than the current action. We argue that these signals represent control processes to update target representations. The current study elucidates the behavioral and neural profiles of different target switching control strategies.

## Acknowledgements

This work was supported by Open Research Area Grant 464-13-003 from the Netherlands Organization for Scientific Research and by European Research Council Consolidator Grant ERC-2013-CoG-615423 to C. N. L. Olivers and by Open Research Area grant DFG PO 548/16-1 to S. Pollmann. We would like to thank Renate Blobel-Lüer and Jörg Stadler for technical and practical assistance during data collection and advice during data preprocessing, Michael Hanke and Falko Kaule for assistance during the setup phase of the study, and Tomas Knapen, Gilles de Hollander and especially Brónagh McCoy for valuable discussions about and suggestions for the fMRI analysis and drift diffusion modeling.

## Supplemental Material

### Three-way repeated measures ANOVA on mean saccade latency with transition type, target availability and display variation as factors

We included different display variations including duplicate targets and duplicate distractors to control for number of targets and prevent potential strategies of looking for anything that occurred twice. In a control analysis, we wanted to check whether these different display variations affected the presence and magnitude of switch costs in each target availability condition. For this purpose, we ran a three-way repeated-measures ANOVA with target availability (both targets vs. one target available), transition type (switch vs. repeat) and display variation (target duplicate vs. distractor duplicate) as factors on saccade latency. This yielded significant main effects of transition type (*F*(1,18) = 22.7, *p <* .001, *η^2^* = 0.56), and target availability (*F*(1,18) = 17.6, *p =* .001, *η^2^* = 0.49), and a two-way interaction between target availability and transition type (*F*(1,18) = 15.6, *p* = .001, *η^2^* = 0.46). None of the other effects reached significance: main effect display variation (*F*(1,18) ≈ 0.0, *p* = .95, *η^2^* < 0.001), the two-way interaction between target availability and display variation (*F*(1,18) = 0.41, *p* = .53, *η^2^* = 0.02), the two-way interaction between transition type and display variation (*F*(1,18) = 0.41, *p* = .53, *η^2^* = 0.02), and three-way interaction (*F*(1,18) = 0.67, *p* = .42, *η^2^* = 0.04). A Bayes Factor analysis confirmed this pattern by showing that the model including the main effects transition type and target availability and the interaction between them explains the data best (*BF* = 6.7 x 10^14^) and was 5.9 times as likely as the next best model that additionally included the main effect display variation. Taken together, these findings suggest that switch costs, expressed in saccade latency, did not depend on specific the display variations.

We ran the same three-way ANOVA on accuracy. This ANOVA indicated that the main effects (target availability: *F*(1,18) = 41.4, *p <* .001, *η^2^* = 0.70; transition type: *F*(1,18) = 35.9, *p <* .001, *η^2^* = 0.67; display variation: *F*(1,18) = 8.91, *p* = .008, *η^2^* = 0.33), and the two-way interactions between target availability and trial transition (*F*(1,18) = 27.3, *p <* .001, *η^2^* = 0.60) and between target availability and display variation (*F*(1,18) = 12.3, *p* = .003, *η^2^* = 0.41) were significant. Neither the two-way interaction between transition type and display variation (*F*(1,18) = 3.5, *p* = .08, *η^2^* = 0.16), nor the three-way interaction (*F*(1,18) = 1.98, *p* = .18, *η^2^* = 0.1) reached significance. Even though accuracy systematically varied with the display variation, such that displays with more targets were associated with higher accuracy, the critical interaction of trial transition and target availability was not affected. Thus, switch costs did not depend on the particular display variation we used.

**Table S1.**
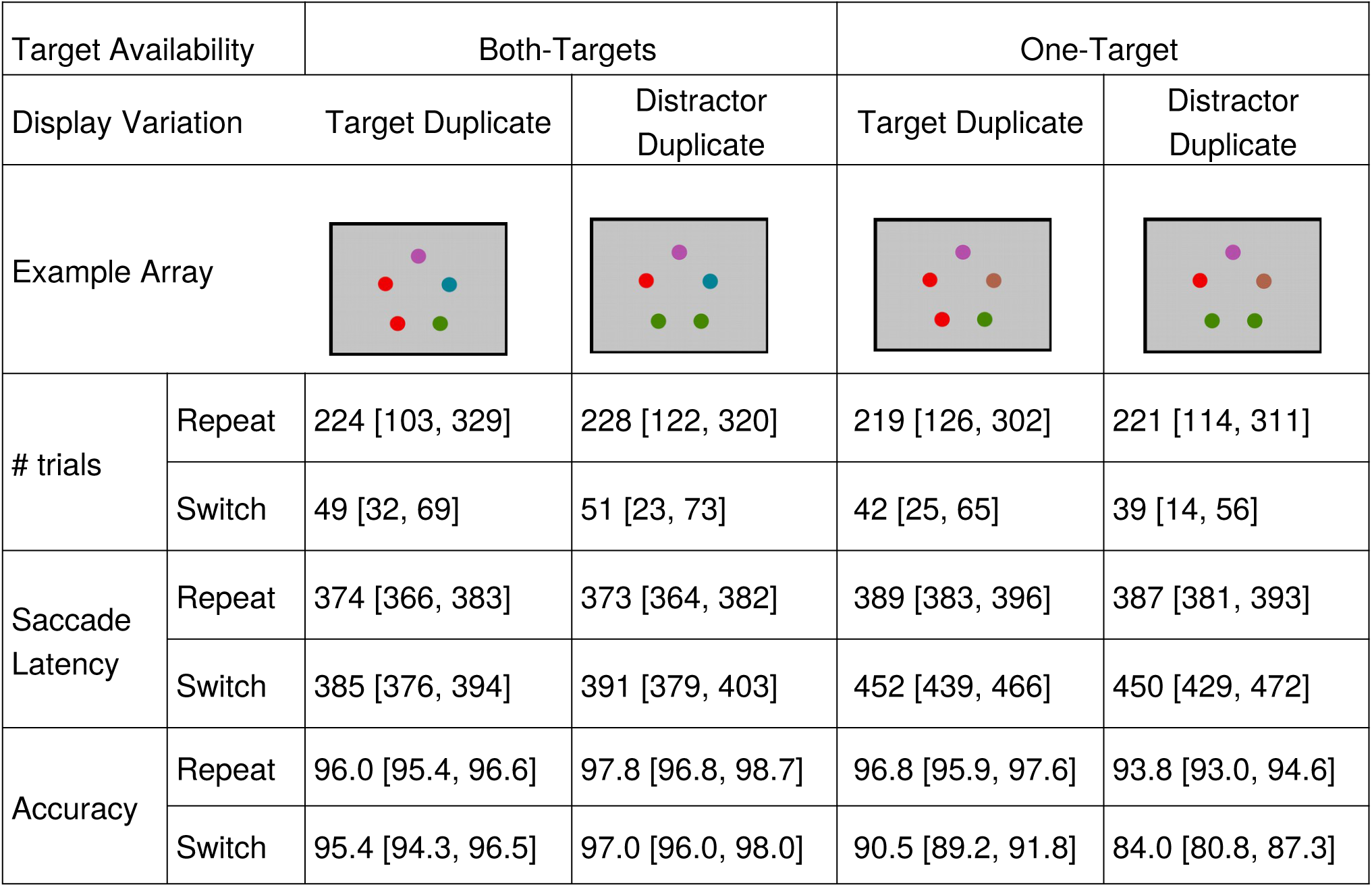
Mean saccade latencies and within-subject 95% CI (Morey, 2008) in ms and percentage correctly fixated targets with within-subject 95% CI for all display variations. Both target availability conditions (both-targets available and one-target available) had two variations of displays, to control for number of target items and duplicate colors in the display. The row *Example Array* depicts an example of each display variation. The row *# trials* represents the mean number (and the min-max range across participants in brackets) of trials per condition. The rows *Repeat* and *Switch* and provide the mean saccade latencies for target switches and target repetitions, for each of these display variations. Examples here are for trials on which red and blue were cued as target colors.

| Target Availability |  | Both-Targets |  | One-Target |  |
| --- | --- | --- | --- | --- | --- |
| Display Variation |  | Target Duplicate | Distractor Duplicate | Target Duplicate | Distractor Duplicate |
| Example Array |  |  |  |  |  |
| # trials | Repeat | 224 [103, 329] | 228 [122, 320] | 219 [126, 302] | 221 [114, 311] |
|  | Switch | 49 [32, 69] | 51 [23, 73] | 42 [25, 65] | 39 [14, 56] |
| Saccade Latency | Repeat | 374 [366, 383] | 373 [364, 382] | 389 [383, 396] | 387 [381, 393] |
|  | Switch | 385 [376, 394] | 391 [379, 403] | 452 [439, 466] | 450 [429, 472] |
| Accuracy | Repeat | 96.0 [95.4, 96.6] | 97.8 [96.8, 98.7] | 96.8 [95.9, 97.6] | 93.8 [93.0, 94.6] |
|  | Switch | 95.4 [94.3, 96.5] | 97.0 [96.0, 98.0] | 90.5 [89.2, 91.8] | 84.0 [80.8, 87.3] |

### Additional Behavioral Analysis reveal no major strategic difference across participants

To gain more insight into different strategies participants might use, we analysed choice behavior in more detail. Overall, these additional analyses suggest that most individuals apply similar strategies when selecting the target to fixate. Specifically, we ran following analyses: (1) Does the streak length (of successive repeat trials) per block vary across participants? For example, if a participant chose the strategy to switch every 6 trials, the streak length should be around 6 with a small standard deviation. In contrast, if a participant switches on each of the first 6 trials of a block and then sticks with one color for the rest of the block, streak length should be smaller and have a higher variability. As Figure S1A demonstrates, most participants used a rather regular switching approach with not too much variability (M = 4.12 repeats, min-max range over participants: 1.99 – 7.52). Exceptions are participant 1 and (to some extent) participant 21 who seem to have a bias toward short streak lengths, and participant 4 who seems to behave rather randomly.

(2) Next, to quantify the regularity with which participants switched targets throughout a block, we split blocks into three parts (first 14, middle 14 and last 13 trials) and checked how many switches fell in each tercile. Figure S1B shows that approximately the same number of switches occurred in each tercile, further supporting the conclusion that participants spread out the switching over the course of a block. Again a handful of participants showed more switches in the first tercile than in the other two, indicative of a strategy that relies on performing the required number of switch trials in the beginning of a block and staying with the same color for the rest of it. We computed the ratio of number of switches in the first tercile with number of switches in the last tercile and correlated this with switch rate, streak length, switch costs in one-target block, switch costs in both-targets blocks, and the difference between these two types of blocks. Importantly, only the correlation of streak length and the ratio of early vs. late switches was significant (Spearman’s ρ (18) = −0.51, p = 0.03), suggesting that participants that switched a lot early in the block, also have shorter streak lengths altogether. This confirms that these individuals most likely applied the strategy of first reaching the required number of switches before sticking with one color for the rest of the block. None of the other correlations was significant (Spearman’s p < 0.26, p > 0.27. Here, we also correlated switch rate across participants with difference of switch costs between both-targets available and one-target available condition, but also this correlation was not significant (Spearman’s ρ (18) = 0.17, p = 0.47), suggesting that the main findings did not depend on how often participants switched between the targets.

(3) Next, we investigated whether the number of targets in the display influenced the likelihood that participant would switch. However, this factor had no influence on target selection (*t*(18) = 0.72, *p* = .48, Cohen’s d = 0.12, BF = 0.3).

(4) To see whether switch costs depended on which color was selected on a trial, we computed switch cost separately for each target color. This analysis revealed no significant effect of color on the switch costs in neither of the target availability conditions (see Figure S1C). Note, when splitting up the trials based on the target color, only 25 observations are left in the smallest cells on average. Therefore, these analyses have to be interpreted with care.

(5) Finally, we checked whether the likelihood to switch depended on the target color by running an ANOVA on switch rate, but also this was not the case, suggesting that participants did not have a preference for switching to a specific color (*F*(3.06,55.13) = 0.75, *p* = 0.53).

**Fig S1.**
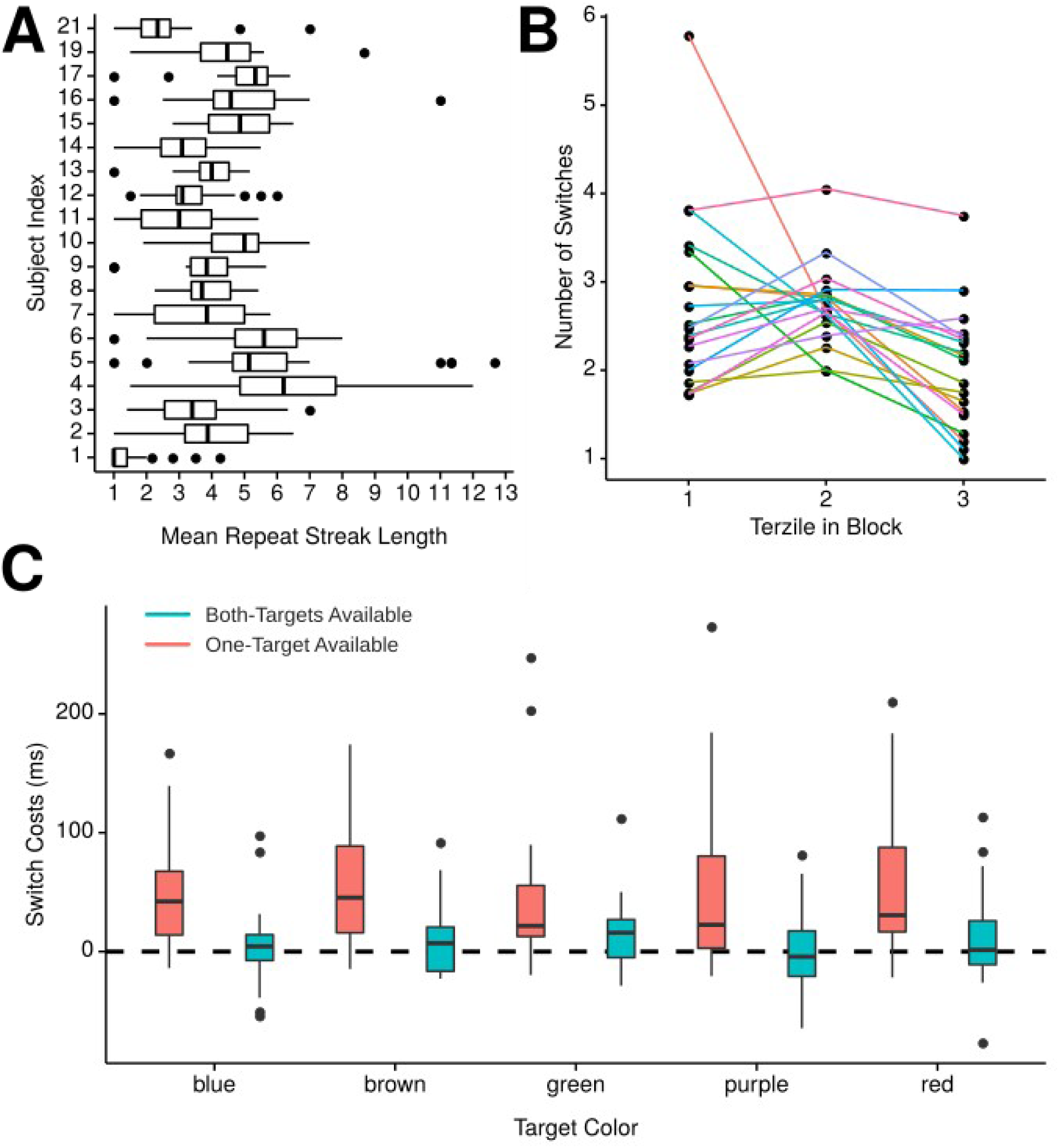
Additional behavioral analysis to investigate strategic difference across participants with respect to switch behavior. (A) Boxplot showing the mean streak length (number of repeat trials in between two successive switch trials). The vertical lines in the box plots represent quartiles. The horizontal line represents the minimum (lower quartile − 1.5 * interquartile range) and maximum (upper quartile + 1.5 * interquartile range) while single dots beyond that range indicate individual outliers. (B) Number of switches in the first, second and third tercile of a block, separately for participants (separate lines). (C) Switch costs for each color in the experiment, separately for each target availability condition.

**Table S2.** All models that were compared during the drift diffusion modeling. The column *Parameters depending on Target Availability* indicates which parameters (*a* and/or *v*), were allowed to vary across the two levels of target availability. The column *Parameters depending on Transition Type* indicates the same, but for transition type. *DIC* represents the deviance information criterion, which can be used to compare models. A lower DIC is better. For all models, we used informative priors (Wiecki et al., 2013), a fixed non-decision time, starting point, and fixed inter-trial variability. All models were checked for convergence.

| Parameters depending on<br>Target availability | Parameters depending on<br>Transition Type | Priors | DIC |
| --- | --- | --- | --- |
| - | - | informative | -50192 |
| $v$ | $v$ | informative | -51323 |
| $a$ | - | informative | -50557 |
| $a, v$ | $v$ | informative | -51486 |
| $a, v$ | $v$ | noninformative | -51486 |

**Table S3.**
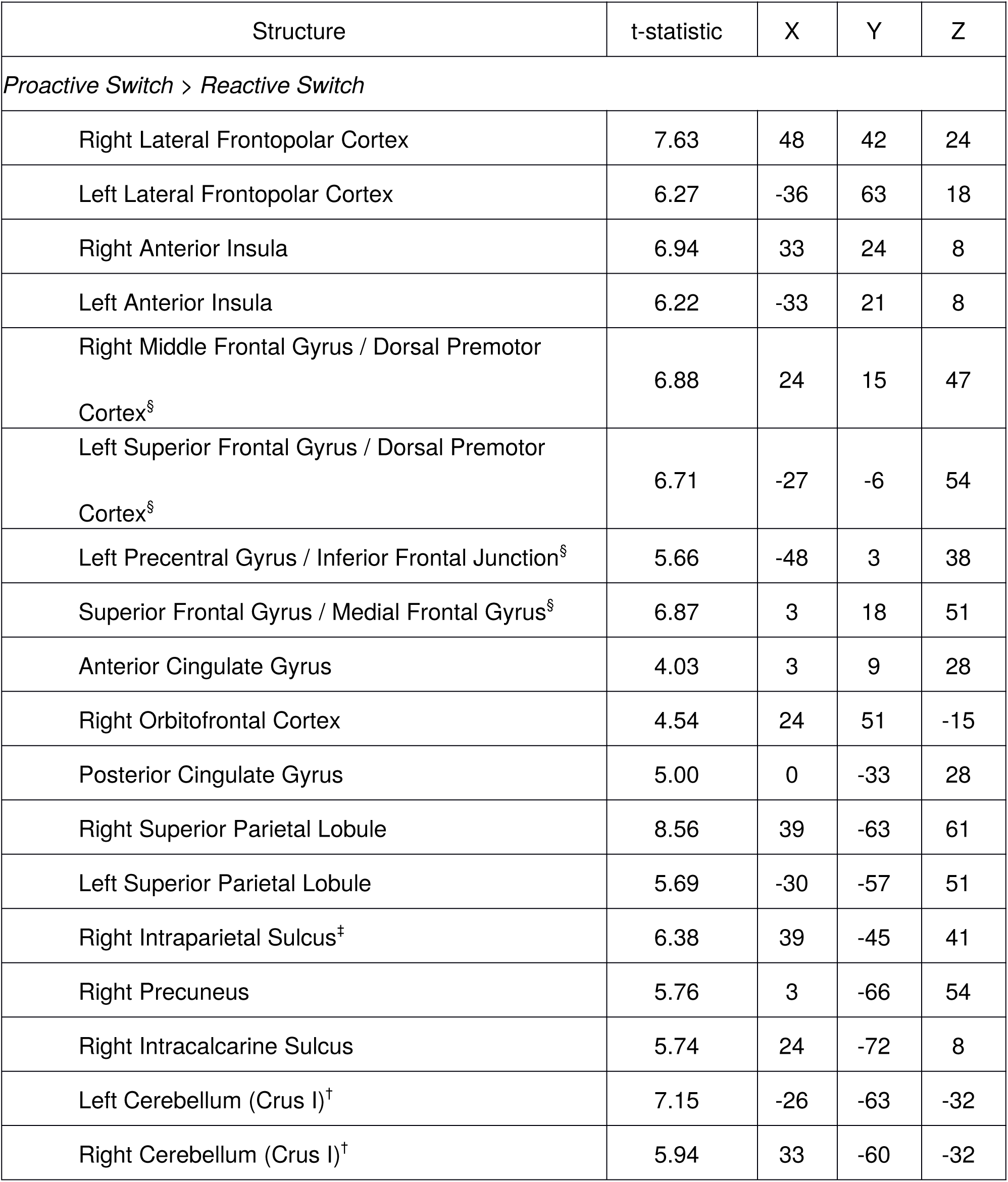
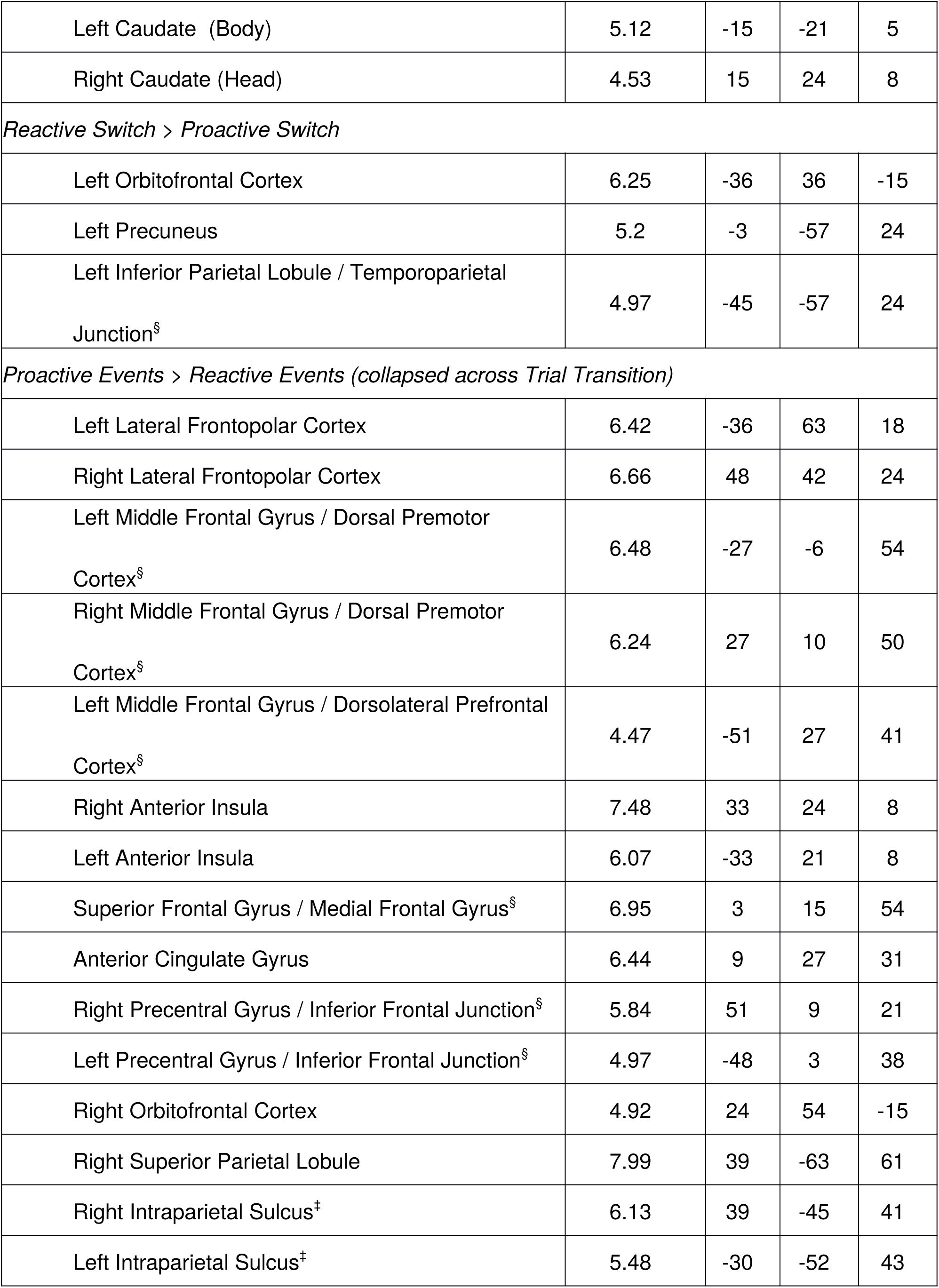
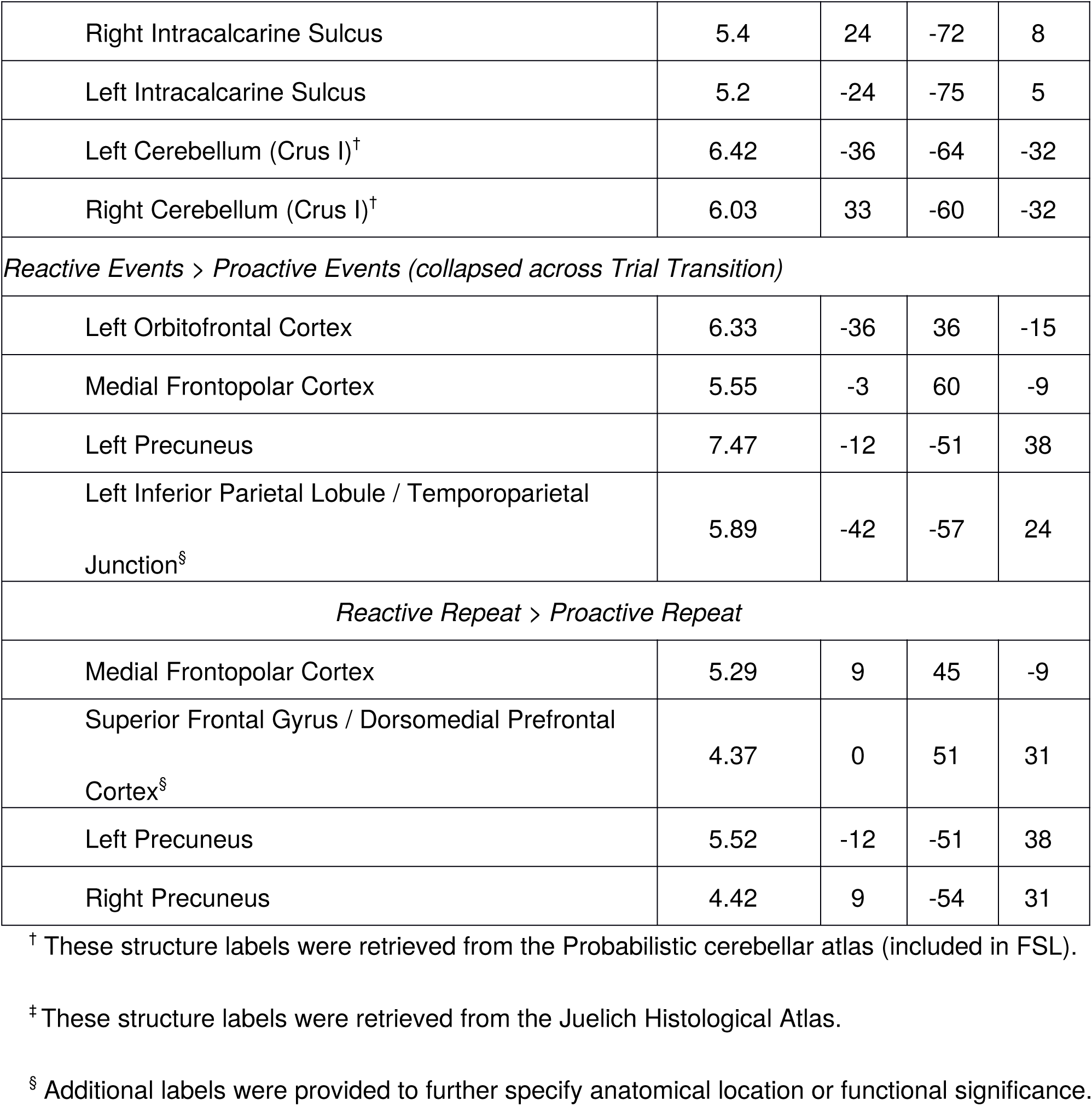
Localization of activations for additionally analyzed contrasts. Coordinates of local maxima are reported in MNI152-space. Large clusters were split into subclusters based on anatomical considerations. Structure labels are based on the Harvard-Oxford anatomical atlas.

| Structure | t-statistic | X | Y | Z |
| --- | --- | --- | --- | --- |
| *Proactive Switch > Reactive Switch* |  |  |  |  |
| Right Lateral Frontopolar Cortex | 7.63 | 48 | 42 | 24 |
| Left Lateral Frontopolar Cortex | 6.27 | -36 | 63 | 18 |
| Right Anterior Insula | 6.94 | 33 | 24 | 8 |
| Left Anterior Insula | 6.22 | -33 | 21 | 8 |
| Right Middle Frontal Gyrus / Dorsal Premotor Cortex<sup>§</sup> | 6.88 | 24 | 15 | 47 |
| Left Superior Frontal Gyrus / Dorsal Premotor Cortex<sup>§</sup> | 6.71 | -27 | -6 | 54 |
| Left Precentral Gyrus / Inferior Frontal Junction<sup>§</sup> | 5.66 | -48 | 3 | 38 |
| Superior Frontal Gyrus / Medial Frontal Gyrus<sup>§</sup> | 6.87 | 3 | 18 | 51 |
| Anterior Cingulate Gyrus | 4.03 | 3 | 9 | 28 |
| Right Orbitofrontal Cortex | 4.54 | 24 | 51 | -15 |
| Posterior Cingulate Gyrus | 5.00 | 0 | -33 | 28 |
| Right Superior Parietal Lobule | 8.56 | 39 | -63 | 61 |
| Left Superior Parietal Lobule | 5.69 | -30 | -57 | 51 |
| Right Intraparietal Sulcus<sup>‡</sup> | 6.38 | 39 | -45 | 41 |
| Right Precuneus | 5.76 | 3 | -66 | 54 |
| Right Intracalcarine Sulcus | 5.74 | 24 | -72 | 8 |
| Left Cerebellum (Crus I)<sup>†</sup> | 7.15 | -26 | -63 | -32 |
| Right Cerebellum (Crus I)<sup>†</sup> | 5.94 | 33 | -60 | -32 |
| Running head: PROACTIVE AND REACTIVE TARGET SELECTION |  |  |  |  |
| Left Caudate (Body) | 5.12 | -15 | -21 | 5 |
| Right Caudate (Head) | 4.53 | 15 | 24 | 8 |
| *Reactive Switch > Proactive Switch* |  |  |  |  |
| Left Orbitofrontal Cortex | 6.25 | -36 | 36 | -15 |
| Left Precuneus | 5.2 | -3 | -57 | 24 |
| Left Inferior Parietal Lobule / Temporoparietal Junction<sup>§</sup> | 4.97 | -45 | -57 | 24 |
| *Proactive Events > Reactive Events (collapsed across Trial Transition)* |  |  |  |  |
| Left Lateral Frontopolar Cortex | 6.42 | -36 | 63 | 18 |
| Right Lateral Frontopolar Cortex | 6.66 | 48 | 42 | 24 |
| Left Middle Frontal Gyrus / Dorsal Premotor Cortex<sup>§</sup> | 6.48 | -27 | -6 | 54 |
| Right Middle Frontal Gyrus / Dorsal Premotor Cortex<sup>§</sup> | 6.24 | 27 | 10 | 50 |
| Left Middle Frontal Gyrus / Dorsolateral Prefrontal Cortex<sup>§</sup> | 4.47 | -51 | 27 | 41 |
| Right Anterior Insula | 7.48 | 33 | 24 | 8 |
| Left Anterior Insula | 6.07 | -33 | 21 | 8 |
| Superior Frontal Gyrus / Medial Frontal Gyrus<sup>§</sup> | 6.95 | 3 | 15 | 54 |
| Anterior Cingulate Gyrus | 6.44 | 9 | 27 | 31 |
| Right Precentral Gyrus / Inferior Frontal Junction<sup>§</sup> | 5.84 | 51 | 9 | 21 |
| Left Precentral Gyrus / Inferior Frontal Junction<sup>§</sup> | 4.97 | -48 | 3 | 38 |
| Right Orbitofrontal Cortex | 4.92 | 24 | 54 | -15 |
| Right Superior Parietal Lobule | 7.99 | 39 | -63 | 61 |
| Right Intraparietal Sulcus<sup>‡</sup> | 6.13 | 39 | -45 | 41 |
| Left Intraparietal Sulcus<sup>‡</sup> | 5.48 | -30 | -52 | 43 |
| Right Intracalcarine Sulcus | 5.4 | 24 | -72 | 8 |
| Left Intracalcarine Sulcus | 5.2 | -24 | -75 | 5 |
| Left Cerebellum (Crus I)<sup>†</sup> | 6.42 | -36 | -64 | -32 |
| Right Cerebellum (Crus I)<sup>†</sup> | 6.03 | 33 | -60 | -32 |
| *Reactive Events > Proactive Events (collapsed across Trial Transition)* |  |  |  |  |
| Left Orbitofrontal Cortex | 6.33 | -36 | 36 | -15 |
| Medial Frontopolar Cortex | 5.55 | -3 | 60 | -9 |
| Left Precuneus | 7.47 | -12 | -51 | 38 |
| Left Inferior Parietal Lobule / Temporoparietal Junction<sup>§</sup> | 5.89 | -42 | -57 | 24 |
| *Reactive Repeat > Proactive Repeat* |  |  |  |  |
| Medial Frontopolar Cortex | 5.29 | 9 | 45 | -9 |
| Superior Frontal Gyrus / Dorsomedial Prefrontal Cortex<sup>§</sup> | 4.37 | 0 | 51 | 31 |
| Left Precuneus | 5.52 | -12 | -51 | 38 |
| Right Precuneus | 4.42 | 9 | -54 | 31 |

<sup>†</sup> These structure labels were retrieved from the Probabilistic cerebellar atlas (included in FSL).
<sup>‡</sup> These structure labels were retrieved from the Juelich Histological Atlas.
<sup>§</sup> Additional labels were provided to further specify anatomical location or functional significance.

**Figure S2.**
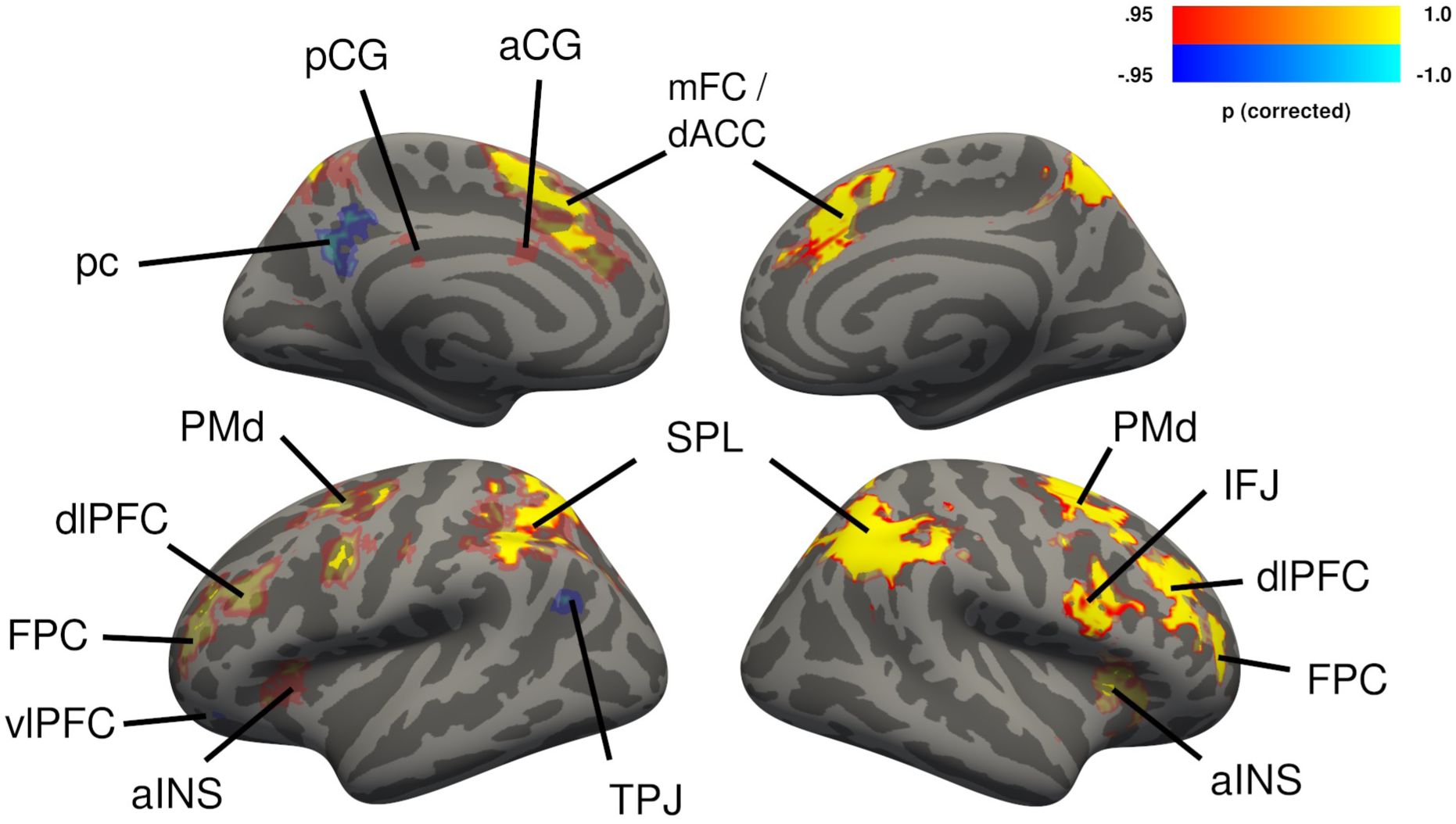
Cerebral activations for free and imposed switches, not taking respective repeat trials into account. Activations shown in yellow-red represent the contrast free switch > imposed switch. Activations shown in blue represent the contrast imposed switch > free switch. Group-level *t*-statistics maps were computed with the tfce-method (S. M. Smith & Nichols, 2009) and corrected for multiple comparisons using nonparametric permutation testing. The resulting P-value maps were thresholded at α = .05 and projected onto the fsaverage surface using registration fusion (Wu et al., 2018), with translucent coloring. In addition, regions that were also significant at α = .01 are shown in saturated colors. Free switches were associated with higher activity than imposed switches across both hemispheres in dorsolateral prefrontal cortex (dlPFC), frontopolar cortex (FPC), dorsal premotor cortex (PMd), right inferior frontal junction (IFJ), anterior insula/frontal operculum cortex (aINS), left posterior cingulate gyrus (pCG), left anterior cingulate gyrus (aCG), bilateral medial frontal cortex/ dorsal anterior cingulate cortex (mFC/dACC) and bilateral superior parietal lobule (SPL). Imposed switches were associated with higher activity than free switches in the right precuneus (pc), left temporoparietal junction and ventrolateral prefrontal cortex (vlPFC).

**Figure S3.**
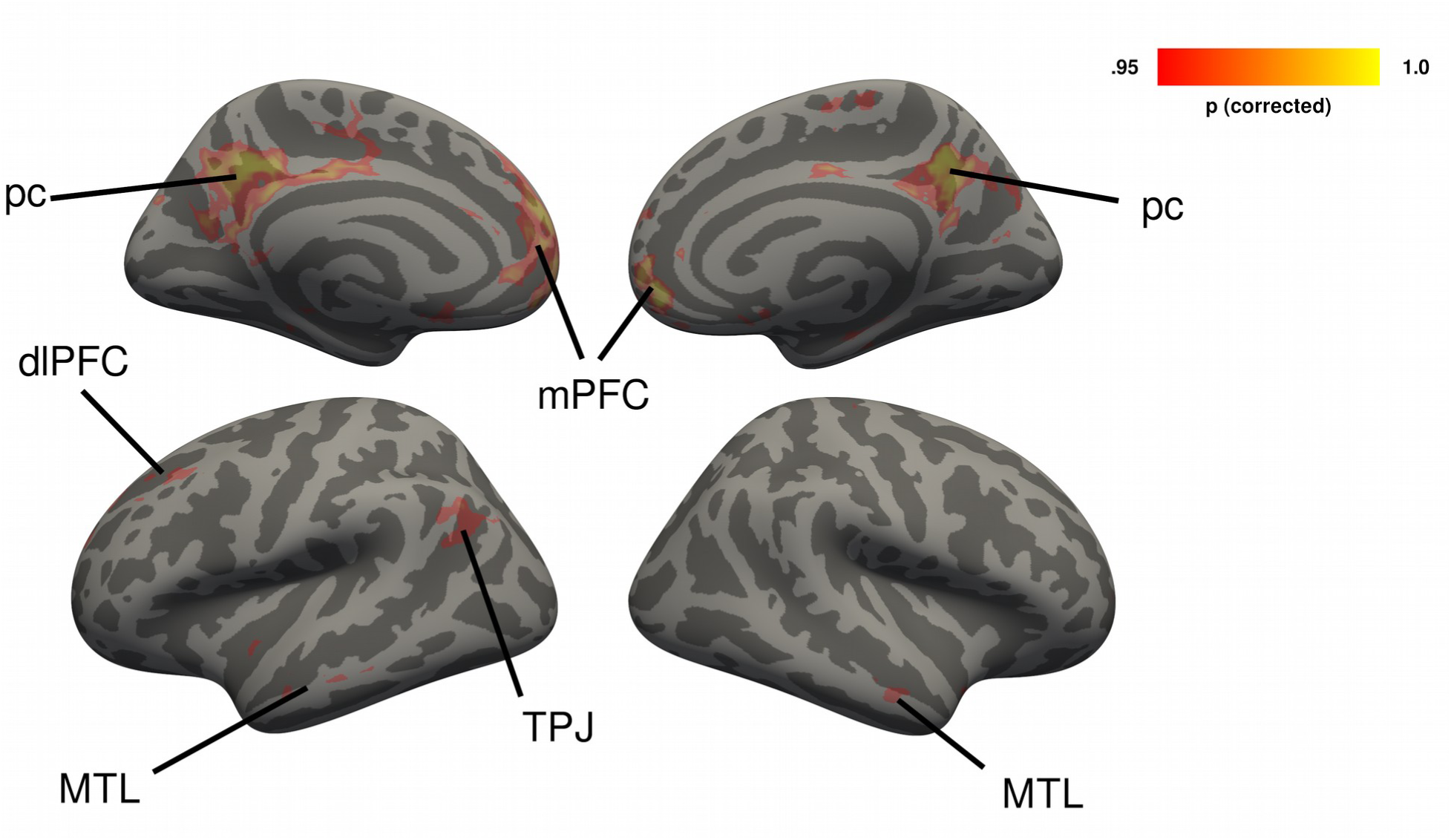
Cerebral regions in which imposed repeats yielded stronger activity than free repeats, not taking respective repeat trials into account (shown in yellow-red). Group-level *t*-statistics maps were computed with the tfce-method (S. M. Smith & Nichols, 2009) and corrected for multiple comparisons using nonparametric permutation testing. The resulting P-value maps were thresholded at α = .05 and projected onto the fsaverage surface using registration fusion (Wu et al., 2018), with translucent coloring. In addition, regions that were also significant at α = .01 are shown in saturated colors. Imposed switches were associated with higher activity than imposed switches across both hemispheres in the precuneus (pc), medial prefrontal cortex (mPFC), medial temporal gyrus (MTG), and the left temperoparietal junction.

**Figure S4.**
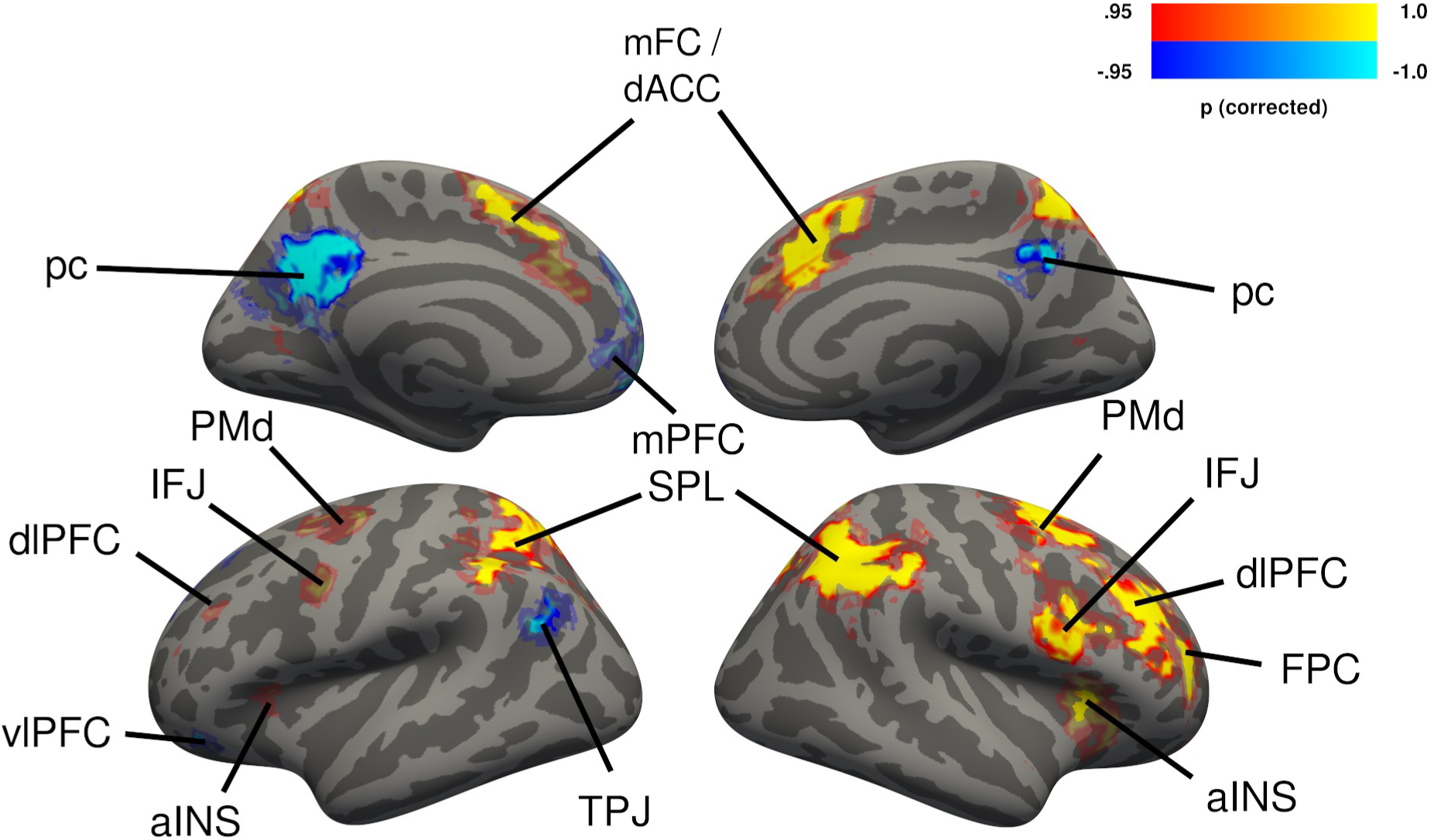
Cerebral activations for the free choice condition compared to the imposed choice condition, irrespective of transition type. Activations shown in yellow-red represent the contrast free events > imposed events. Activations shown in blue represent the contrast imposed events > free events. Group-level *t*-statistics maps were computed with the tfce-method (S. M. Smith & Nichols, 2009) and corrected for multiple comparisons using nonparametric permutation testing. The resulting P-value maps were thresholded at α = .05 and projected onto the fsaverage surface using registration fusion (Wu et al., 2018), with translucent coloring. In addition, regions that were also significant at α = .01 are shown in saturated colors. Free events were associated with higher activity than imposed events across both hemispheres in dorsolateral prefrontal cortex (dlPFC), frontopolar cortex (FPC), dorsal premotor cortex (PMd), right inferior frontal junction (IFJ), anterior insula/frontal operculum cortex (aINS), bilateral medial frontal cortex/ dorsal anterior cingulate cortex (mFC/dACC) and bilateral superior parietal lobule (SPL). Imposed events were associated with higher activity than free events in the precuneus (pc), left temporoparietal junction, medial prefrontal cortex (mPFC), ventrolateral prefrontal cortex (vlPFC).

## References

Abraham, A., Pedregosa, F., Eickenberg, M., Gervais, P., Muller, A., Kossaifi, J., … Varoquaux, G. (2014). Machine Learning for Neuroimaging with Scikit-Learn. Frontiers in Neuroinformatics, 8(February), 1–10. http://doi.org/10.3389/fninf.2014.00014

Arrington, C. M., & Logan, G. D. (2004). The cost of a voluntary task switch. Psychological Science, 15(9), 610–615. http://doi.org/10.1111/j.0956-7976.2004.00728.x

Arrington, C. M., & Logan, G. D. (2005). Voluntary task switching: Chasing the elusive homunculus. Journal of Experimental Psychology: Learning Memory and Cognition, 31(4), 683–702. http://doi.org/10.1037/0278-7393.31.4.683

Avants, B. B., Epstein, C. L., Grossman, M., & Gee, J. C. (2008). Symmetric diffeomorphic image registration with cross-correlation: Evaluating automated labeling of elderly and neurodegenerative brain. Medical Image Analysis, 12 (1), 26–41. http://doi.org/10.1016/j.media.2007.06.004

Badre, D., & D’Esposito, M. (2009). Is the rostro-caudal axis of the frontal lobe hierarchical? Nature Reviews Neuroscience, 10(9), 659–669. http://doi.org/10.1038/nrn2667

Barrett, D. J. K., & Zobay, O. (2014). Attentional control via parallel target-templates in dual-target search. PLoS ONE, 9(1), 1–9. http://doi.org/10.1371/journal.pone.0086848

Behzadi, Y., Restom, K., Liau, J., & Liu, T. T. (2007). A component based noise correction method (CompCor) for BOLD and perfusion based fMRI. NeuroImage. 37(1), 90–101. http://doi.org/10.1016/j.neuroimage.2007.04.042

Botvinick, M., Nystrom, L. E., Fissell, K., Carter, C. S., & Cohen, J. D. (1999). Conflict monitoring versus selection-for-action in anterior cingulate cortex. Nature, 402(11), 179–181. http://doi.org/10.1038/46035

Brass, M., & von Cramon, D. Y. (2004). Decomposing Components of Task Preparation with Functional Magnetic Resonance Imaging. Journal of Cognitive Neuroscience, 16(4), 609–620. http://doi.org/10.1162/089892904323057335

Braver, T. S. (2012). The variable nature of cognitive control: A dual mechanisms framework. Trends in Cognitive Sciences, 16(2), 106–113. http://doi.org/10.1016/j.tics.2011.12.010

Braver, T. S., Cohen, J. D., Nystrom, L. E., Jonides, J., Smith, E. E., & Noll, D. C. (1997). A parametric study of prefrontal cortex involvement in human working memory. NeuroImage. 5 (1), 49–62. http://doi.org/10.1006/nimg.1996.0247

Braver, T. S., Paxton, J. L., Locke, H. S., & Barch, D. M. (2009). Flexible neural mechanisms of cognitive control within human prefrontal cortex. Proceedings of the National Academy of Sciences of the United States of America, 106(18), 7351–7356. http://doi.org/10.1073/pnas.0808187106

Braver, T. S., Reynolds, J. R., & Donaldson, D. I. (2003). Neural mechanisms of transient and sustained cognitive control during task switching. Neuron, 39(4), 713–726. http://doi.org/10.1016/S0896-6273(03)00466-5

Bunge, S. A., Kahn, I., Wallis, J. D., Miller, E. K., & Wagner, A. D. (2003). Neural Circuits Subserving the Retrieval and Maintenance of Abstract Rules. Journal of Neurophysiology, 90(5), 3419–3428. http://doi.org/10.1152/jn.00910.2002

Bunge, S. A., Ochsner, K. N., Desmond, J. E., Glover, G. H., & Gabrieli, J. D. E. (2001). Prefrontal regions involved in keeping information in and out of mind. Brain, 124, 2074–2086. http://doi.org/10.1093/brain/124.10.2074

Burgess, G. C., & Braver, T. S. (2010). Neural mechanisms of interference control in working memory: Effects of interference expectancy and fluid intelligence. PLoS ONE, 5(9), 1–11. http://doi.org/10.1371/journal.pone.0012861

Cavanagh, J. F., & Frank, M. J. (2014). Frontal theta as a mechanism for cognitive control. Trends in Cognitive Sciences.,18 (8), 414–421. http://doi.org/10.1016/j.tics.2014.04.012

Cohen, M. X. (2014). A neural microcircuit for cognitive conflict detection and signaling. Trends in Neurosciences, 37 (9), 480–490. http://doi.org/10.1016/j.tins.2014.06.004

Cole, M. W., & Schneider, W. (2007). The cognitive control network: Integrated cortical regions with dissociable functions. NeuroImage, 37(1), 343–360. http://doi.org/10.1016/j.neuroimage.2007.03.071

Dale, A. M., Fischl, B., & Sereno, M. I. (1999). Cortical surface-based analysis: I. Segmentation and surface reconstruction. NeuroImage 9(2), 179–194. http://doi.org/10.1006/nimg.1998.0395

Dalmaijer, E. S., Mathôt, S., & Van der Stigchel, S. (2013). PyGaze: An open-source, cross-platform toolbox for minimal-effort programming of eyetracking experiments. Behavior Research Methods, 46, 913–921. http://doi.org/10.3758/s13428-013-0422-2

de Hollander, G., & Knapen, T. (2018). VU-Cog-Sci/nideconv: First alpha version nideconv (Version v0.1.0). Zenodo. http://doi.org/http://doi.org/10.5281/zenodo.1463839

De Pisapia, N., & Braver, T. S. (2006). A model of dual control mechanisms through anterior cingulate and prefrontal cortex interactions. Neurocomputing, 69(10–12), 1322–1326. http://doi.org/10.1016/j.neucom.2005.12.100

Demanet, J., De Baene, W., Arrington, C. M., & Brass, M. (2013). Biasing free choices: The role of the rostral cingulate zone in intentional control. NeuroImage, 72, 207–213. http://doi.org/10.1016/j.neuroimage.2013.01.052

Desimone, R., & Duncan, J. (1995). Neural Mechanisms of Selective Visual Attention. Annual Review of Neuroscience, 18(1), 193–222. http://doi.org/10.1146/annurev.neuro.18.1.193

Dombrowe, I., Donk, M., & Olivers, C. N. L. (2011). The costs of switching attentional sets. Attention, Perception, & Psychophysics, 73, 2481–2488. http://doi.org/10.3758/s13414-011-0198-3

Donner, T. H., Siegel, M., Fries, P., & Engel, A. K. (2009). Buildup of Choice-Predictive Activity in Human Motor Cortex during Perceptual Decision Making. Current Biology, 19(18), 1581–1585. http://doi.org/10.1016/j.cub.2009.07.066

Dosenbach, N. U. F., Fair, D. A., Cohen, A. L., Schlaggar, B. L., & Petersen, S. E. (2008). A dual-networks architecture of top-down control. Trends in Cognitive Sciences, 12(3), 99–105. http://doi.org/10.1016/j.tics.2008.01.001

Dosenbach, N. U. F., Visscher, K. M., Palmer, E. D., Miezin, F. M., Wenger, K. K., Kang, H. C., … Petersen, S. E. (2006). A Core System for the Implementation of Task Sets. Neuron, 50(5), 799–812. http://doi.org/10.1016/j.neuron.2006.04.031

Dove, A., Pollmann, S., Schubert, T., Wiggins, C. J., & Yves von Cramon, D. (2000). Prefrontal cortex activation in task switching: An event-related fMRI study. Cognitive Brain Research, 9(1), 103–109. http://doi.org/10.1016/S0926-6410(99)00029-4

Duncan, J. (2010). The multiple-demand (MD) system of the primate brain: mental programs for intelligent behaviour. Trends in Cognitive Sciences, 14(4), 172–179. http://doi.org/10.1016/j.tics.2010.01.004

Duprez, J., Gulbinaite, R., & Cohen, M. X. (2018). Midfrontal theta phase coordinates behaviorally relevant brain computations during response conflict. BioRxiv, 1–38. https://doi.org/10.1101/502716

Elton, A., & Gao, W. (2015). Task-positive Functional Connectivity of the Default Mode Network Transcends Task Domain. Journal of Cognitive Neuroscience, 27(12), 2369–2381. http://doi.org/10.1162/jocn

Esteban, O., Markiewicz, C. J., Blair, R. W., Moodie, C. A., Isik, A. I., Erramuzpe, A., … Gorgolewski, K. J. (2018). FMRIPrep: a robust preprocessing pipeline for functional MRI. Nature Methods, 1–10. http://doi.org/10.1038/s41592-018-0235-4

Fedorenko, E., Duncan, J., & Kanwisher, N. (2013). Broad domain generality in focal regions of frontal and parietal cortex. Proceedings of the National Academy of Sciences, 110(41), 16616–16621. http://doi.org/10.1073/pnas.1315235110

Fonov, V., Evans, A., McKinstry, R., Almli, C., & Collins, D. (2009). Unbiased nonlinear average age-appropriate brain templates from birth to adulthood. NeuroImage, 47 (Supplement 1), S102. http://doi.org/10.1016/S1053-8119(09)70884-5

Forstmann, B. U., Brass, M., Koch, I., & von Cramon, D. Y. (2006). Voluntary selection of task sets revealed by functional magnetic resonance imaging. Journal of Cognitive Neuroscience, 18(3), 388–398. http://doi.org/10.1162/jocn.2006.18.3.388

Found, A., & Müller, H. J. (1996). Searching for unknown feature targets on more than one dimension: investigating a “dimension-weighting” account. Perception & Psychophysics, 58(1), 88–101. http://doi.org/10.3758/BF03205479

Gmeindl, L., Chiu, Y. C., Esterman, M. S., Greenberg, A. S., Courtney, S. M., & Yantis, S. (2016). Tracking the will to attend: Cortical activity indexes self-generated, voluntary shifts of attention. Attention, Perception, and Psychophysics, 78(7), 2176–2184. http://doi.org/10.3758/s13414-016-1159-7

Gorgolewski, K. J., Burns, C. D., Madison, C., Clark, D., Halchenko, Y. O., Waskom, M. L., & Ghosh, S. S. (2011). Nipype: A Flexible, Lightweight and Extensible Neuroimaging Data Processing Framework in Python. Frontiers in Neuroinformatics, 5(August). http://doi.org/10.3389/fninf.2011.00013

Gorgolewski, K. J., Esteban, O., Markiewicz, C. J., Ziegler, E., Ellis, D. G., Jarecka, D., … Ghosh, S. (2018). nipy/nipype: 1.1.6. http://doi.org/10.5281/ZENODO.1560596

Gramfort, A., Luessi, M., Larson, E., Engemann, D. a., Strohmeier, D., Brodbeck, C., … Hämäläinen, M. (2013). MEG and EEG data analysis with MNE-Python. Frontiers in Neuroscience, 7(December), 1–13. http://doi.org/10.3389/fnins.2013.00267

Greenberg, A. S., Esterman, M., Wilson, D., Serences, J. T., & Yantis, S. (2010). Control of spatial and feature-based attention in frontoparietal cortex. The Journal of Neuroscience: The Official Journal of the Society for Neuroscience, 30(43), 14330–14339. http://doi.org/10.1523/JNEUROSCI.4248-09.2010

Greve, D. N., & Fischl, B. (2009). Accurate and robust brain image alignment using boundary-based registration. NeuroImage, 48(1), 63–72. http://doi.org/10.1016/j.neuroimage.2009.06.060

Hopfinger, J. B., Buonocore, M. H., & Mangun, G. R. (2000). The neural mechanisms of top-down attentional control. Nature Neuroscience, 3(3), 284–291. http://doi.org/10.1038/72999

Huang, L., & Pashler, H. (2007). A Boolean map theory of visual attention. Psychological Review, 114(3), 599–631. http://doi.org/10.1037/0033-295X.114.3.599

Huntenburg, J. M. (2014). Evaluating nonlinear coregistration of BOLD EPI and T1w images. Doctoral Dissertation, Free University Berlin.

Irlbacher, K., Kraft, A., Kehrer, S., & Brandt, S. A. (2014). Mechanisms and neuronal networks involved in reactive and proactive cognitive control of interference in working memory. Neuroscience and Biobehavioral Reviews, 46, 58–70. http://doi.org/10.1016/j.neubiorev.2014.06.014

Ito, S., Stuphorn, V., Brown, J. W., & Schall, J. D. (2003). Performance monitoring by the anterior cingulate cortex during saccade countermanding. Science 302 (5642), 120–122. http://doi.org/10.1126/science.1087847

Jenkinson, M., Bannister, P., Brady, M., & Smith, S. M. (2002). Improved optimization for the robust and accurate linear registration and motion correction of brain images. NeuroImage. 17 (2), 825–841. http://doi.org/10.1016/S1053-8119(02)91132-8

Jiang, J., Beck, J., Heller, K., & Egner, T. (2015). An insula-frontostriatal network mediates flexible cognitive control by adaptively predicting changing control demands. Nature Communications, 6(May), 1–11. http://doi.org/10.1038/ncomms9165

Jiang, J., Wagner, A. D., & Egner, T. (2018). Integrated externally and internally generated task predictions Jointly guide cognitive control in prefrontal cortex. ELife, 7, 1–23. http://doi.org/10.7554/eLife.39497

Juola, J. F., Botella, J., & Palacios, A. (2004). Task- and location-switching effects on visual attention. Perception & Psychophysics, 66(8), 1303–17. http://doi.org/10.3758/BF03195000

Karayanidis, F., Mansfield, E. L., Galloway, K. L., Smith, J. L., Provost, A., & Heathcote, A. (2009). Anticipatory reconfiguration elicited by fully and partially informative cues that validly predict a switch in task. *Cognitive*, Affective and Behavioral Neuroscience. 9 (2), 202–215. http://doi.org/10.3758/CABN.9.2.202

Kerns, J. G., Cohen, J. D., MacDonald, A. W., Cho, R. Y., Stenger, V. A., & Carter, C. S. (2004). Anterior Cingulate Conflict Monitoring and Adjustments in Control. Science, 303 (5660), 1023–1026. http://doi.org/10.1126/science.1089910

Kim, C., Cilles, S. E., Johnson, N. F., & Gold, B. T. (2012). Domain general and domain preferential brain regions associated with different types of task switching: A Meta-Analysis. Human Brain Mapping, 33(1), 130–142. http://doi.org/10.1002/hbm.21199

Klein, A., Ghosh, S. S., Bao, F. S., Giard, J., Häme, Y., Stavsky, E., … Keshavan, A. (2017). Mindboggling morphometry of human brains. PLoS Computational Biology, 13 (2), e1005350. http://doi.org/10.1371/journal.pcbi.1005350

Kloosterman, N. A., de Gee, J. W., Werkle-Bergner, M., Lindenberger, U., Garrett, D. D., & Fahrenfort, J. J. (2019). Humans strategically shift decision bias by flexibly adjusting sensory evidence accumulation. eLife, 8, e37321. https://doi.org/10.7554/eLife.3732

Konishi, M., McLaren, D. G., Engen, H., & Smallwood, J. (2015). Shaped by the past: The default mode network supports cognition that is independent of immediate perceptual input. PLoS ONE, 10(6), 1–18. http://doi.org/10.1371/journal.pone.0132209

Liesefeld, H. R. (2018) Estimating the Timing of Cognitive Operations With MEG/EEG Latency Measures: A Primer, a Brief Tutorial, and an Implementation of Various Methods, Frontiers in Neuroscience, 12(October), 1–11. doi: 10.3389/fnins.2018.00765

Liston, C., Matalon, S., Hare, T. A., Davidson, M. C., & Casey, B. J. (2006). Anterior Cingulate and Posterior Parietal Cortices Are Sensitive to Dissociable Forms of Conflict in a Task-Switching Paradigm. Neuron, 50(4), 643–653. http://doi.org/10.1016/j.neuron.2006.04.015

Liu, T., & Jigo, M. (2017). Limits in feature-based attention to multiple colors. Attention, Perception, & Psychophysics, 79, 2327–2337. http://doi.org/10.3758/s13414-017-1390-x

Liu, T., Slotnick, S. D., Serences, J. T., & Yantis, S. (2003). Cortical Mechanisms of Feature-based Attentional Control. Cerebral Cortex, 13(12), 1334–1343. http://doi.org/10.1093/cercor/bhg080

Luck, S. J. (2014). An introduction to the event-related potential technique (2nd ed.). Cambridge, MA: MIT press.

Maljkovic, V., & Nakayama, K. (1994). Priming of pop-out: I. Role of features. Memory & Cognition, 22(6), 657–672. http://doi.org/10.3758/BF03209251

Mansouri, F. A., Koechlin, E., Rosa, M. G. P., & Buckley, M. J. (2017). Managing competing goals — a key role for the frontopolar cortex. Nature Neuroscience Reviews, 18(11), 645–657. http://doi.org/10.1038/nrn.2017.111

Marklund, P., & Persson, J. (2012). Context-dependent switching between proactive and reactive working memory control mechanisms in the right inferior frontal gyrus. NeuroImage, 63(3), 1552–1560. http://doi.org/10.1016/j.neuroimage.2012.08.016

Mathôt, S., Schreij, D., & Theeuwes, J. (2012). OpenSesame: An open-source, graphical experiment builder for the social sciences. Behavior Research Methods, 44(2), 314–324. http://doi.org/10.3758/s13428-011-0168-7

Meiran, N. (1996). Reconfiguration of processing mode prior to task performance. Journal of Experimental Psychology: Learning, Memory, and Cognition, 22(6), 1423–1442. http://doi.org/10.1037/0278-7393.22.6.1423

Meiran, N. (2010). Task switching and cognitive self control. In Self control in Society, Mind and Brain (pp. 202–220). http://doi.org/10.1093/acprof:oso/9780195391381.001.0001

Menneer, T., Barrett, D. J. K., Phillips, L., Donnelly, N., & Cave, K. R. (2007). Costs in Searching for Two Targets: Dividing Search Across Target Types Could Improve Airport Security Screening. Applied Cognitive Psychology, (21), 915–932. http://doi.org/10.1002/acp.1305

Menon, V., & Uddin, L. Q. (2010). Saliency, switching, attention and control: a network model of insula function. Brain Structure & Function, 214(5–6), 655–667. http://doi.org/10.1007/s00429-010-0262-0

Miller, E. K., & Cohen, J. D. (2001). An integrative theory of prefrontal cortex function. Annual Review of Neuroscience, 24, 167–202. https://doi.org/10.1146/annurev.neuro.24.1.167

Miller, J., Patterson, T., & Ulrich, R. (1998). Jackknife-based method for measuring LRP onset latency differences. Psychophysiology, 35(1), 99–115. doi: 10.1111/1469-8986.3510099

Moore, K. S., & Weissman, D. H. (2010). Involuntary transfer of a top-down attentional set into the focus of attention: Evidence from a contingent attentional capture paradigm. Attention, Perception, & Psychophysics, 72(6), 1495–1509. https://doi.org/10.3758/APP.72.6.1495

Morey, R. D. (2008). Confidence Intervals from Normalized Data: A correction to Cousineau (2005). Tutorials in Quantitative Methods for Psychology, 4(2), 61–64. http://doi.org/10.3758/s13414-012-0291-2

Niendam, T. A., Laird, A. R., Ray, K. L., Dean, Y. M., Glahn, D. C., & Carter, C. S. (2012). Meta-analytic evidence for a superordinate cognitive control network subserving diverse executive functions. Cognitive, Affective and Behavioral Neuroscience, 12(2), 241–268. http://doi.org/10.3758/s13415-011-0083-5

Oberauer, K. (2002). Access to information in working memory: exploring the focus of attention. Journal of Experimental Psychology. Learning, Memory, and Cognition, 28(3), 411–421. http://doi.org/10.1037/0278-7393.28.3.411

Olivers, C. N. L., & Eimer, M. (2011). On the difference between working memory and attentional set. Neuropsychologia, 49(6), 1553–1558. http://doi.org/10.1016/j.neuropsychologia.2010.11.033

Olivers, C. N. L., Peters, J., Houtkamp, R., & Roelfsema, P. R. (2011). Different states in visual working memory: When it guides attention and when it does not. Trends in Cognitive Sciences, 15(7), 327–334. http://doi.org/10.1016/j.tics.2011.05.004

Orr, J. M., & Banich, M. T. (2013). The Neural Mechanisms Underlying Internally and Externally Guided Task Selection. Neuroimage, 80309(303), 1–30. http://doi.org/10.1016/j.neuroimage.2013.08.047

Ort, E., Fahrenfort, J. J., & Olivers, C. N. L. (2017). Lack of Free Choice Reveals the Cost of Having to Look for More Than One Object. Psychological Science, 28(8), 1137–1147. http://doi.org/10.1177/0956797617705667

Ort, E., Fahrenfort, J. J., & Olivers, C. N. L. (2018). Lack of free choice reveals the cost of multiple-target search within and across feature dimensions. Attention, Perception, and Psychophysics, (80), 1904–1917. http://doi.org/10.3758/s13414-018-1579-7

Ort E., Fahrenfort, J. J., ten Cate, T., and Olivers, C. N. L. (2019) Humans can efficiently look for, but not select multiple visual objects, bioRxiv. doi: http://dx.doi.org/10.1101/653030v1

Passingham, R. E., Bengtsson, S. L., & Lau, H. C. (2010). Medial frontal cortex: from self-generated action to reflection on one’s own performance. Trends in Cognitive Sciences, 14(1), 16–21. http://doi.org/10.1016/j.tics.2009.11.001

Poldrack, R. A., Mumford, J. A., & Nichols, T. E. (2011). Handbook of functional MRI data analysis. Cambridge: Cambridge University Press. http://doi.org/https://doi.org/10.1017/CBO9780511895029

Pollmann, S. (2016). Frontopolar Resource Allocation in Human and Nonhuman Primates. Trends in Cognitive Sciences, 20(2), 84–86. http://doi.org/10.1016/j.tics.2015.11.006

Pollmann, S., Weidner, R., Müller, H. J., Maertens, M., & von Cramon, D. Y. (2006). Selective and interactive neural correlates of visual dimension changes and response changes. NeuroImage, 30(1), 254–265. http://doi.org/10.1016/j.neuroimage.2005.09.013

Pollmann, S., Weidner, R., Müller, H. J., & von Cramon, D. Y. (2000). A fronto-posterior network involved in visual dimension changes. Journal of Cognitive Neuroscience, 12(3), 480–494. http://doi.org/10.1162/089892900562156

Power, J. D., Mitra, A., Laumann, T. O., Snyder, A. Z., Schlaggar, B. L., & Petersen, S. E. (2014). Methods to detect, characterize, and remove motion artifact in resting state fMRI. NeuroImage, 84, 320–341. http://doi.org/10.1016/j.neuroimage.2013.08.048

Power, J. D., & Petersen, S. E. (2013). Control-related systems in the human brain. Current Opinion in Neurobiology, 23(2), 223–228. http://doi.org/10.1016/j.conb.2012.12.009

Raichle, M. E. (2015). The Brain’s Default Mode Network. Annual Review of Neuroscience, 38(1), 433–447. http://doi.org/10.1146/annurev-neuro-071013-014030

Raichle, M. E., Macleod, A. M., Snyder, A. Z., Powers, W. J., Gusnard, D. A., & Shulman, G. L. (2001). A default mode of brain function. Proceedings of the National Academy of Sciences, 98(2), 676–682. http://doi.org/10.1073/pnas.98.2.676

Ratcliff, R., & Rouder, J. N. (1998). Modeling Response Times for Two-Choice Decisions. Psychological Science, 9(5), 347–356. http://doi.org/10.1111/1467-9280.00067

Ryman, S. G., El Shaikh, A. A., Shaff, N. A., Hanlon, F. M., Dodd, A. B., Wertz, C. J., … Mayer, A. R. (2018). Proactive and reactive cognitive control rely on flexible use of the ventrolateral prefrontal cortex. Human Brain Mapping, (January), 1–12. http://doi.org/10.1002/hbm.24424

Rypma, B., & D’Esposito, M. (1999). The roles of prefrontal brain regions in components of working memory: Effects of memory load and individual differences. Proceedings of the National Academy of Sciences, 96 (11), 6558–6563. http://doi.org/10.1073/pnas.96.11.6558

Schmitz, F., & Voss, A. (2012). Decomposing task-switching costs with the diffusion model. Journal of Experimental Psychology: Human Perception and Performance, 38(1), 222–250. http://doi.org/10.1037/a0026003

Seeley, W. W., Menon, V., Schatzberg, A. F., Keller, J., Glover, G. H., Kenna, H., … Greicius, M. D. (2007). Dissociable Intrinsic Connectivity Networks for Salience Processing and Executive Control. Journal of Neuroscience, 27(9), 2349–2356. http://doi.org/10.1523/JNEUROSCI.5587-06.2007

Shenhav, A., Botvinick, M. M., & Cohen, J. D. (2013). The expected value of control: An integrative theory of anterior cingulate cortex function. Neuron, 79(2), 217–240. http://doi.org/10.1016/j.neuron.2013.07.007

Shulman, G. L. (2002). Two Attentional Processes in the Parietal Lobe. Cerebral Cortex, 12(11), 1124–1131. http://doi.org/10.1093/cercor/12.11.1124

Singmann, H., Bolker, B., Westfall, J., Aust, F., Højsgaard, S., Fox, J., … Love, J. (2016). afex: analysis of factorial experiments. R package version 0.16-1. R Package Version 0.16-1.

Slagter, H. A., Giesbrecht, B., Kok, A., Weissman, D. H., Kenemans, J. L., Woldorff, M. G., & Mangun, G. R. (2007). fMRI evidence for both generalized and specialized components of attentional control. Brain Research, 1177(1), 90–102. http://doi.org/10.1016/j.brainres.2007.07.097

Slagter, H. A., Weissman, D. H., Giesbrecht, B., Kenemans, J. L., Mangun, G. R., Kok, A., & Woldorff, M. G. (2006). Brain regions activated by endogenous preparatory set shifting as revealed by fMRI. Cognitive, Affective and Behavioral Neuroscience, 6(3), 175–189. http://doi.org/10.3758/CABN.6.3.175

Smallwood, J., Tipper, C., Brown, K., Baird, B., Engen, H., Michaels, J. R., … Schooler, J. W. (2013). Escaping the here and now: Evidence for a role of the default mode network in perceptually decoupled thought. NeuroImage, 69, 120–125. http://doi.org/10.1016/j.neuroimage.2012.12.012

Smith, A. B., Taylor, E., Brammer, M., & Rubia, K. (2004). Neural Correlates of Switching Set as Measured in Fast, Event-Related Functional Magnetic Resonance Imaging. Human Brain Mapping, 21(4), 247–256. http://doi.org/10.1002/hbm.20007

Smith, S. M., & Brady, J. M. (1997). SUSAN - A new approach to low level image processing. International Journal of Computer Vision, 23(1), 45–78. http://doi.org/10.1023/A:1007963824710

Smith, S. M., & Nichols, T. E. (2009). Threshold-free cluster enhancement: Addressing problems of smoothing, threshold dependence and localisation in cluster inference. NeuroImage, 44(1), 83–98. http://doi.org/10.1016/j.neuroimage.2008.03.061

Smith, V., Mitchell, D. J., & Duncan, J. (2018). Role of the Default Mode Network in Cognitive Transitions. Cerebral Cortex, 28(October), 3685–3696. http://doi.org/10.1093/cercor/bhy167

Sohn, M.-H., Ursu, S., Anderson, J. R., Stenger, V. A., & Carter, C. S. (2000). The role of prefrontal cortex and posterior parietal cortex in task switching. Proceedings of the National Academy of Sciences, 97(24), 13448–13453. https://doi.org/10.1073/pnas.240460497

Soon, C. S., Brass, M., Heinze, H. J., & Haynes, J. D. (2008). Unconscious determinants of free decisions in the human brain. Nature Neuroscience, 11(5), 543–545. http://doi.org/10.1038/nn.2112

Spitzer, B., & Haegens, S. (2017). Beyond the Status Quo: A Role for Beta Oscillations in Endogenous Content (Re-) Activation. Eneuro, 4(4). http://doi.org/10.1523/ENEURO.0170-17.2017

Spreng, R. N. (2012). The fallacy of a “task-negative” network. Frontiers in Psychology, 3(MAY), 1–5. http://doi.org/10.3389/fpsyg.2012.00145

Spreng, R. N., DuPre, E., Selarka, D., Garcia, J., Gojkovic, S., Mildner, J., … Turner, G. R. (2014). Goal-Congruent Default Network Activity Facilitates Cognitive Control. Journal of Neuroscience, 34(42), 14108–14114. http://doi.org/10.1523/JNEUROSCI.2815-14.2014

Taylor P. C. J., Rushworth, M. F. S., & Nobre, A. C. (2008) Choosing where to attend and the medial frontal cortex: an FMRI study. Journal of Neurophysiology, 100(3):1397–1406. 10.1152/jn.90241.2008

Treiber, J. M., White, N. S., Steed, T. C., Bartsch, H., Holland, D., Farid, N., … Chen, C. C. (2016). Characterization and correction of geometric distortions in 814 Diffusion Weighted Images. PLoS ONE, 11(3), e0152472. http://doi.org/10.1371/journal.pone.0152472

Tustison, N., Avants, B. B., Cook, P. A., Zheng, Y., Egan, A., Yushkevich, P. U., & Gee, J. C. (2010). N4ITK: Improved N3 Bias Correction (Vol. 29). IEEE Transactions of Medical Imaging.

Ullsperger, M., Danielmeier, C., & Jocham, G. (2014). Neurophysiology of Performance Monitoring and Adaptive Behavior. Physiological Reviews, 94(1), 35–79. http://doi.org/10.1152/physrev.00041.2012

van Driel, J., Ort, E., Fahrenfort, J. J., & Olivers, C. N. L. (2019). Beta and theta oscillations differentially support free versus forced control over multiple-target search. Journal of Neuroscience, 149, 114–128. http://doi.org/https://doi.org/10.1523/JNEUROSCI.2547-18.2018

Vatansever, D., Menon, D. K., & Stamatakis, E. A. (2017). Default mode contributions to automated information processing. Proceedings of the National Academy of Sciences, 114(48), 12821–12826. http://doi.org/10.1073/pnas.1710521114

Wager, T. D., Jonides, J., & Reading, S. (2004). Neuroimaging studies of shifting attention: A meta-analysis. NeuroImage, 22(4), 1679–1693. http://doi.org/10.1016/j.neuroimage.2004.03.052

Wang, S., Peterson, D. J., Gatenby, J. C., Li, W., Grabowski, T. J., & Madhyastha, T. M. (2017). Evaluation of Field Map and Nonlinear Registration Methods for Correction of Susceptibility Artifacts in Diffusion MRI. Frontiers in Neuroinformatics, 11, 17. http://doi.org/10.3389/fninf.2017.00017

Wiecki, T. V., Sofer, I., & Frank, M. J. (2013). HDDM: Hierarchical Bayesian estimation of the Drift-Diffusion Model in Python. Frontiers in Neuroinformatics, 7(August), 1–10. http://doi.org/10.3389/fninf.2013.00014

Winkler, A. M., Ridgway, G. R., Webster, M. A., Smith, S. M., & Nichols, T. E. (2014). Permutation inference for the general linear model. NeuroImage, 92, 381–397. http://doi.org/10.1016/j.neuroimage.2014.01.060

Wisniewski, D., Goschke, T., & Haynes, J. D. (2016). Similar coding of freely chosen and externally cued intentions in a fronto-parietal network. NeuroImage, 134, 450–458. http://doi.org/10.1016/j.neuroimage.2016.04.044

Wisniewski, D., Reverberi, C., Tusche, A., & Haynes, J. D. (2015). The neural representation of voluntary task-set selection in dynamic environments. Cerebral Cortex, 25(12), 4715–4726. http://doi.org/10.1093/cercor/bhu155

Woolrich, M. W., Ripley, B. D., Brady, M., & Smith, S. M. (2001). Temporal autocorrelation in univariate linear modeling of FMRI data. NeuroImage, 14(6), 1370–1386. http://doi.org/10.1006/nimg.2001.0931

Wu, J., Ngo, G. H., Greve, D., Li, J., He, T., Fischl, B., … Yeo, B. T. T. (2018). Accurate nonlinear mapping between MNI volumetric and FreeSurfer surface coordinate systems. Human Brain Mapping, 39(9), 3793–3808. http://doi.org/10.1002/hbm.24213

Zhang, J., Kriegeskorte, N., Carlin, J. D., & Rowe, J. B. (2013). Choosing the Rules: Distinct and Overlapping Frontoparietal Representations of Task Rules for Perceptual Decisions. Journal of Neuroscience, 33(29), 11852–11862. http://doi.org/10.1523/JNEUROSCI.5193-12.2013

Zhang, Y., Brady, M., & Smith, S. M. (2001). Segmentation of brain MR images through a hidden Markov random field model and the expectation-maximization algorithm. IEEE Transactions on Medical Imaging, 20(1), 45–57. http://doi.org/10.1109/42.906424

